# Silicon-rhodamine-enabled Identification (SeeID) for Near-Infrared Light Controlled Proximity Labeling In Vitro and In Vivo

**DOI:** 10.1101/2024.12.19.627432

**Authors:** Wenjing Wang, Hongyang Guo, Xiaosa Yan, Xuanzhen Pan, Xiaofei Wang, Yiming Rong, Zexiao Bai, Liwan Zhang, Zhaofa Wu, Xinyu Zhao, Weiren Huang, Wei Qin, Ling Chu

**Author notes:** These authors contributed equally to this work (Wenjing Wang, Hongyang Guo, Xiaosa Yan).

## Abstract

Advancement in fluorescence imaging techniques enables the study of protein dynamics and localization with unprecedented spatiotemporal resolution. However, current imaging tools are unable to elucidate dynamic protein interactomes underlying imaging observations. In contrast, proteomics tools such as proximity labeling enable the analysis of protein interactomes at a single time point but lack information about protein dynamics. We herein developed Silicon-rhodamine-enabled Identification (SeeID) for near-infrared light controlled proximity labeling that could bridge the gap between imaging and proximity labeling. SeeID was benchmarked through characterization of various organelle-specific proteomes and the KRAS protein interactome. The fluorogenic nature of SiR allows for intracellular proximity labeling with high subcellular specificity. Leveraging SiR as both a fluorophore and a photocatalyst, we developed a protocol that allows the study of dynamic protein interactomes of Parkin during mitophagy. We discovered the association of the proteasome complex with Parkin at early time points, indicating the involvement of the ubiquitin-proteasome system for protein degradation in the early phase of mitophagy. In addition, by virtue of the deep tissue penetration of near-infrared light, we achieved spatiotemporally controlled proximity labeling in vivo across the mouse brain cortex with a labeling depth of ∼2 mm using an off-the-shelf 660 nm LED light set-up.

## Main

Live-cell fluorescence imaging has revolutionized how protein localization and dynamics are studied. Proteins execute their functions by interacting with a complex network of biomolecules. However, such interactome information is typically absent in fluorescence imaging analyses. Although fluorescence tagging followed by immunoprecipitation-mass spectrometry (IP-MS) allows the mapping of protein localization and interactions in human cells, IP-MS is unable to detect transient or weak protein-protein interactions (PPIs)^1^.

Proximity labeling (PL), coupled with mass spectrometry, has emerged as a powerful approach to study PPIs^2–5^. PL labels proximal biomolecules with a covalent handle, enabling the capture of transient and weak PPIs. The early development of PL techniques involved fusing the protein of interest (POI) to a peroxidase^6–11^, biotin ligase^11–17^ or Pup ligase^18^, facilitating the installation of biotin handles on neighboring biomolecules. These tagged molecules are then isolated and analyzed by MS to map the POI interactome. Light-activated PL offers complementary methods for investigating PPIs with spatiotemporal precision. These methods leverage genetically encoded^19–22^ or synthetic photocatalysts^23–34^ to locally generate highly reactive intermediates that label proximal proteins. The tagged protein residues are then analyzed by MS. However, the non-fluorescent nature of these PL tags necessitates immunostaining to analyze the spatial distribution of the POI, precluding real-time tracking^35, 36^. Moreover, spatiotemporally controlled PL in vivo is still challenging^5, 19, 36–38^.

Silicon Rhodamine (SiR) and its derivatives are a class of fluorogenic near-infrared (NIR)-emitting fluorophores^39–45^. The fluorogenicity of SiR originates from the equilibrium between its non-fluorescent spirocyclic form and fluorescent zwitterionic form. SiR becomes fluorescent when conjugated to self-labeling tags (e.g., Halo-Tag, SNAP-tag) but remains non-fluorescent in hydrophobic environments or aqueous solutions. Herein, we developed SeeID (<u>S</u>ilicon-rhodamin<u>e</u>-<u>e</u>nabled <u>Id</u>entification) for NIR-light controlled proximity labeling (PL) in vitro and in vivo. The fluorescence nature of SiR supports the development of a live-cell imaging-guided PL protocol to simultaneously track and map POI interactomes. This method leverages the fluorogenic properties of SiR to achieve intracellular PL with high spatial specificity. More importantly, the NIR-activated system enables spatiotemporally controlled PL in vivo.

## Results

We began by repurposing SiR as a photocatalyst. Visible-light photocatalysts exert their catalytic ability through light-induced electron transfer (EnT) or energy transfer (ET). Inspired by the pioneering works of Macmillan^24, 26, 30, 31^ and Rovis^28^, which demonstrated NIR-light activated PL through EnT-mediated nitrene generation. The ability of SiR to generate a nitrene intermediate from perfluorinated azide (**1**) upon photoactivation (Extended Data Fig. 1a) was first investigated. With NADH as the reductant, perfluorinated aniline (PFAA, **2**) was obtained in 51% yields (Extended Data Fig.1b, Supplementary information Fig. S1), confirming nitrene intermediate formation. However, when we extended this protocol to the labeling of BSA using PFAA-biotin, no labeling of BSA was detected by western blot (Extended Data Fig. 1c), likely due to quenching of the light-excited SiR by molecular oxygen.

We then turned our attention to a possible ET pathway. In photodynamic therapy (PDT), light-induced ET between the photosensitizer and molecular oxygen has been explored to generate singlet oxygen for therapeutic purposes^46^. This ET pathway has similarly been applied in the development of light-activated PL strategies^20–22, 35, 36, 47, 48^. SiR has been used as a photocatalyst for in situ generation of tetrazines via photooxidation^49, 50^. However, it remains uncertain whether SiR, optimized for fluorescence imaging, can generate sufficient singlet oxygen for PL. To investigate, 10 μM SiR-CA (Fig. 1a) was reacted with 10 μM Halo protein, forming a covalent SiR-CA-Halo complex that turned on SiR fluorescence (Extended Data Fig. 2a). Addition of biotinylated perfluorinated aniline probe B1 (Extended Data Fig. 2b), followed by irradiation with NIR light (660 nm) resulted in Halo protein biotinylation confirmed by western blot analysis. Substitution of B1 with biotinylated aniline BA increased labeling efficiency, whereas alkylamine B2 reduced it (Fig. 1b). The generation of singlet oxygen was verified by adding the singlet oxygen fluorescence sensor SOSG to the reaction^51^. Irradiation with 660 nm LED light increased SOSG fluorescence, indicating singlet oxygen formation (Fig. 1c). The intensity of SOSG fluorescence is consistent with the labeling efficiency (Fig. 1d). The addition of a singlet oxygen quencher, Vitamin C (NaVc), reduced both SOSG fluorescence and labeling efficiency, further confirming singlet oxygen involvement in Halo protein biotinylation (Fig. 1c). Notably, the fluorogenic nature of SiR-CA minimizes singlet oxygen generation upon irradiation of SiR-CA alone (Fig. 1c). To demonstrate that this fluorogenicity reduces nonspecific background labeling, BSA was labeled with either SiR-CA or a non-fluorogenic NIR photosensitizer, Methylene Blue (MB). SiR-CA labeled BSA only in the presence of Halo protein, whereas MB labeled BSA regardless of Halo protein presence (Fig. 1e). Various nucleophilic probes containing alkyne handles (Fig. 1a and Extended Data Fig.2d), enabling click chemistry and subsequent LC-MS/MS analysis, were screened to further improve the labeling efficiency (Extended Data Fig. 2e). All probes but AP achieved efficient labeling (Fig. 1f). While labeling could be detected upon 5 mins light exposure, we observed an increase of labeling band intensity with longer irradiation time (Fig.1g).

**Figure 1.**
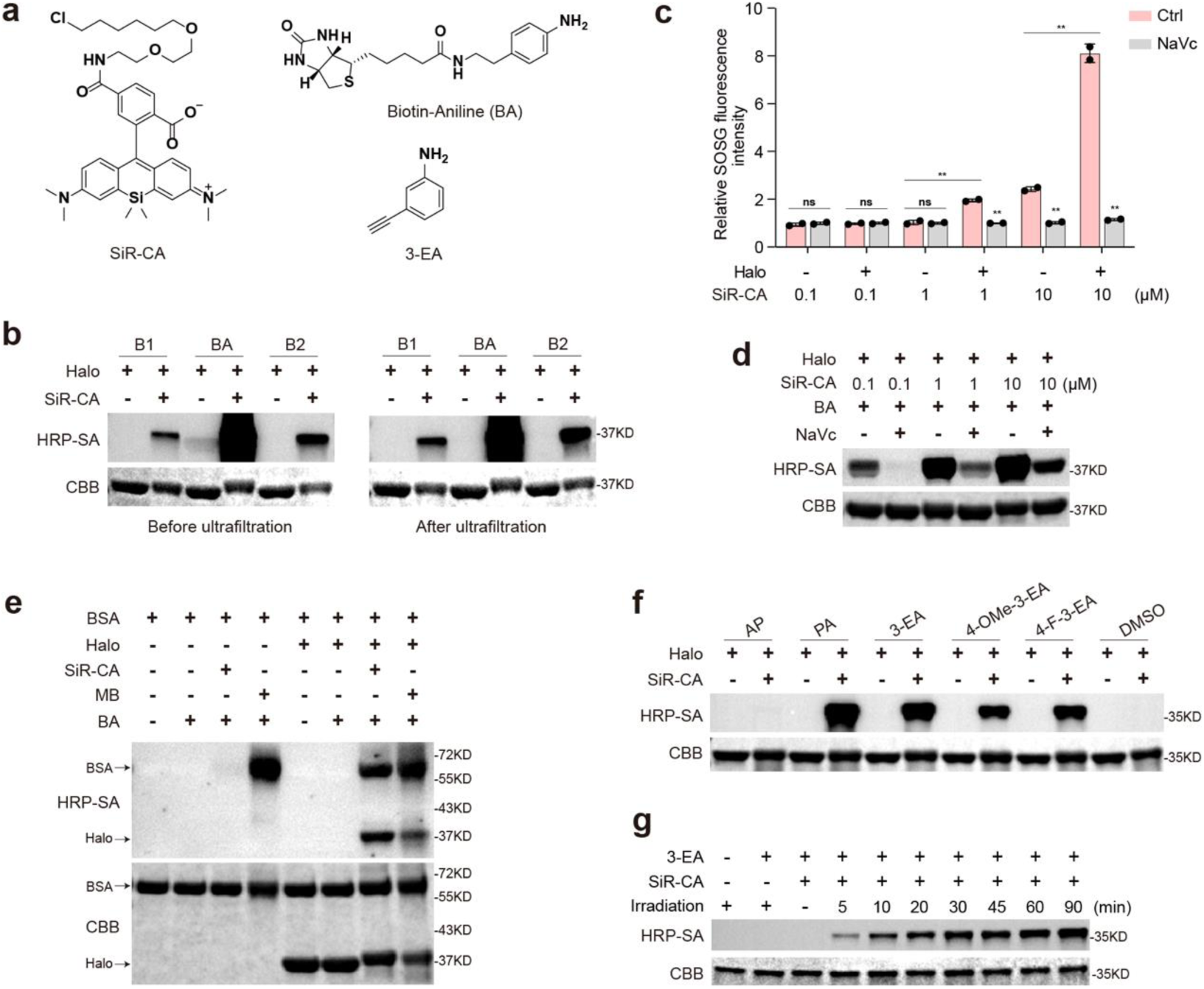
In vitro protein labeling by SeeID. **a,** Chemical structures of photocatalyst SiR-CA, Biotin-conjugated probe BA, and alkynyl probe 3-EA. **b,** 10 μM Halo protein labeling 100 μM by Biotin-conjugated probes with or without 10 μM SiR-CA in 100 μL PBS. The mixtures were irradiated under a 660 nm LED light of 200 mW/cm^2^ intensity at 4 ℃ for 1 hour. Half of the labeled samples were desalted in Ultra Centrifugal Filter (10 kDa MWCO) and recovered with PBS. Biotinylated-Halo was detected by HRP-conjugated streptavidin (HRP-SA) and total proteins were detected by Coomassie Brilliant Blue (CBB) staining. **c-d,** Control experiments demonstrating singlet oxygen generation by SiR-CA only in the presence of Halo protein. SiR-CA mixed with an equivalent amount of Halo protein in PBS, with or without singlet oxygen quencher 5 mM sodium ascorbate (NaVc), was irradiated under 660 nm LED light for 1 hour, followed by singlet oxygen sensor green (SOSG) to detect singlet oxygen production (**c**). Labeling of 10 μM Halo with 100 μM BA using different concentrations of SiR-CA (0.1, 1,10 μM), with or without 5 mM sodium ascorbate (**d**). **e,** 5 μM BSA labeling by 100 μM BA with 10 μM SiR-CA or MB in the presence of 10 μM Halo under a 660 nm LED light of 200 mW/cm^2^ intensity at 4 ℃ for 30 minutes. Biotinylated-Halo and -BSA was detected by HRP-conjugated streptavidin (HRP-SA) and total proteins were detected by CBB staining. **f,** 10 μM Halo protein were labeled by 100 μM alkynyl probes with or without 10 μM SiR-CA in PBS, and desalted in Ultra Centrifugal Filter (10 kDa MWCO), followed by click reaction with Biotin-N_3_ via Cu(I)-catalyzed azide-alkyne cycloaddition (CuAAC). **g,** Evaluation of labeling kinetics. A labeling mixture of 10 μM Halo Protein, 10 μM SiR-CA and 100 μM 3-EA in 1.2 mL PBS were irradiated under 660 nm LED light of 200 mW/cm^2^ intensity at 4 ℃, and 100 μL aliquots were taken from the mixture at the indicated time and analyzed by western blot and CBB staining.

Encouraged by the initial results, labeling conditions in cellulo were optimized. Nucleophilic probes containing alkyne handles validated in vitro were first screened. HEK293T cells transiently expressing Flag-Halo-NLS (Nuclear Localization Signals) were treated with 1 μM SiR-CA and 500 μM nucleophilic probes, then irradiated with 660 nm LED light (50 mW/cm²) for 30 minutes. Following cell lysis, proteins labeled with an alkyne handle underwent Cu(I)-catalyzed azide-alkyne cycloaddition (CuAAC) with Biotin-N_3_ and were analyzed by western blot. Among the tested probes, 3-ethynylaniline (3-EA) exhibited the highest labeling efficiency (Fig. 2a). Thus, 3-EA was selected as the nucleophilic probe for subsequent experiments.

**Figure 2.**
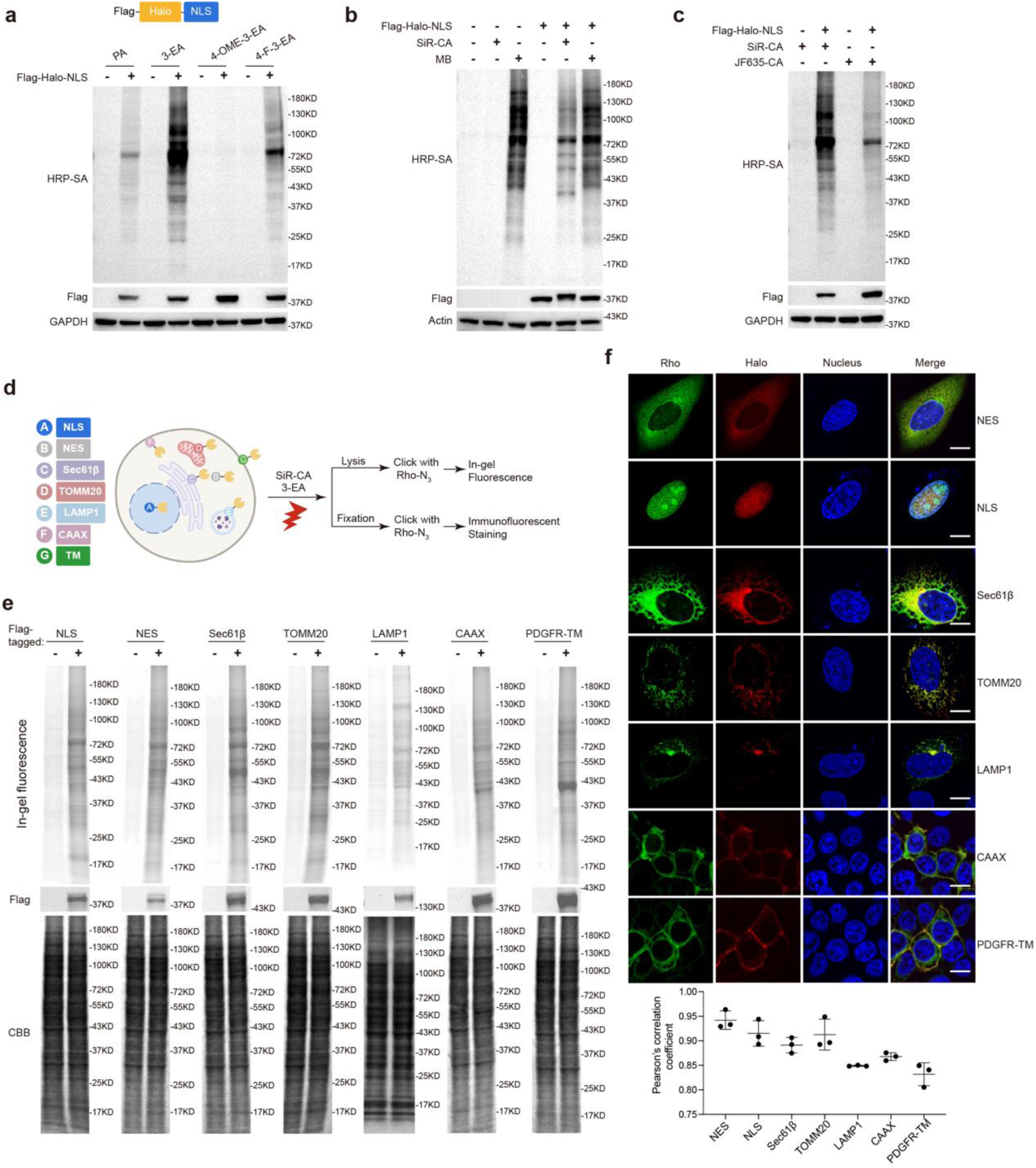
SeeID labeling in living cells. **a,** Evaluation of different labeling probes. HEK293T cells transiently expressing Flag-were treated with 1 μM SiR-CA in HBSS buffer for 1 hour, followed by 500 μM alkynyl probe for 10 minutes. Cells were irradiated under 50 mW/cm^2^ 660 nm LED light at room temperature for 30 minutes, followed by lysing in RIPA, and the lysates were subjected to click reaction with Biotin-N_3_ via CuAAC. After the click reaction, the proteins were extracted using chloroform-methanol extraction and recovered in PBS with 0.5% SDS. The biotinylated proteins and Flag-Halo-NLS expression were analyzed by western blot. **b,** The fluorogenicity of SiR-CA enables Halo-dependent proximal labeling in HEK293T cells. A nonfluorogenic photocatalyst MB was used as a control. HEK293T cells transiently expressing Flag-Halo-NLS were labeled by 500 μM 3-EA using 1 μM SiR-CA or MB. The resulting cell lysates were analyzed by western blot. **c,** Evaluation of different photocatalysts. HEK293T cells transiently expressing Flag-Halo-NLS were labeled by 500 μM 3-EA using 1 μM SiR-CA or JF635-CA. The resulting cell lysates were analyzed by western blot. **d,** Protocols of SeeID labeling of various organelles. **e,** In-gel fluorescence detection of labeled proteins on different organelles in HEK293T cells. **f,** Confocal immunofluorescence images of various organelles labeling in U2OS cells (CAAX and PDGFR-TM were expressed in HEK293T cells). Scale bars: 10 μm. The Pearson’s correlation coefficients were quantified by Image J.

The performance of various NIR photocatalysts were subsequently evaluated. Consistent with prior results, SiR-CA exhibited a Halo protein-dependent labeling pattern (Fig. 2b, lines 2 and 5). The specificity of SiR-CA was preliminarily assessed using confocal microscopy (Extended Data Fig. 3a). Following PL, cells were fixed, clicked with Biotin-N_3_ via click reaction and stained with Streptavidin-FITC to visualize biotinylated proteins. Colocalization (Pearson correlation coefficient (PCC) = 0.98) was observed between the 633 nm channel (Halo-SiR) and the 488 nm channel (Streptavidin-FITC), confirming labeling specificity.

In contrast, photocatalyst MB demonstrated Halo-independent labeling (Fig. 2b, lines 3 and 6), and confocal analysis revealed nonspecific labeling of cytosolic proteins (Fig. 2a and Extended Data Fig.3a). JF635, a derivative of SiR, exhibited reduced labeling efficiency relative to SiR-CA (Fig. 2c and Extended Data Fig. 3a). The cytotoxicity of SiR-CA and MB were assessed using the cell viability assay. SiR-CA exhibited minimal cytotoxicity, whereas MB showed a concentration-dependent cytotoxic effect (Extended Data Fig. 3b).

With the optimal photocatalyst and probe for cellular PL identified, organelle-specific proteomes were characterized using SeeID. U2OS or HEK293T cells transiently expressing Halo-tagged nuclear localization signals (NLS), nuclear export signals (NES), endoplasmic reticulum (ER) marker Sec61β, mitochondrial outer membrane protein TOMM20, lysosomal membrane protein LAMP1, and plasma membrane-targeting motifs (CAAX, PDGFR-TM) were subjected to confocal imaging and the optimized PL protocol (Fig. 2d). Western blot analysis revealed distinct labeling patterns with varying intensities across samples (Fig. 2e). Correct localization of Halo-tagged organelle markers was confirmed by confocal imaging (Fig. 2f). Confocal microscopy demonstrated strong colocalization between labeled proteins (stained with Rhodamine) and Halo-tagged organelle markers (PCC = 0.83-0.94), confirming the high spatial specificity of SeeID across all tested organelles (Fig. 2f).

The performance of SeeID was benchmarked through proteomic profiling of the local proteome at the endoplasmic reticulum (ER) membrane, a widely studied compartment for evaluating the spatial specificity of PL methods (Fig. 3a). HEK293T cells expressing the Halo-Sec61β fusion protein (SeeID-ERM) were treated with 1 μM SiR-CA and 500 μM 3-EA, followed by irradiation with 660 nm LED light at 50 mW/cm² for 30 minutes. A negative control lacking HaloTag expression was included, along with a spatial reference using SeeID-NES to nonspecifically label all cytosolic proteins. Biotinylated proteins were captured with streptavidin beads, digested with trypsin on-beads to release peptides, and labeled with three isotopes (“light,” “medium,” and “heavy”) via dimethyl labeling before pooling and LC-MS/MS analysis. Peptides were identified, and dimethyl labeling ratios were quantified using MaxQuant. Proteins with at least two peptides were quantified based on their median ratios. In total, 1652 proteins were quantified across three biological replicates, which demonstrated high correlation (Extended Data Fig. 4a and Table S1).

**Figure 3.**
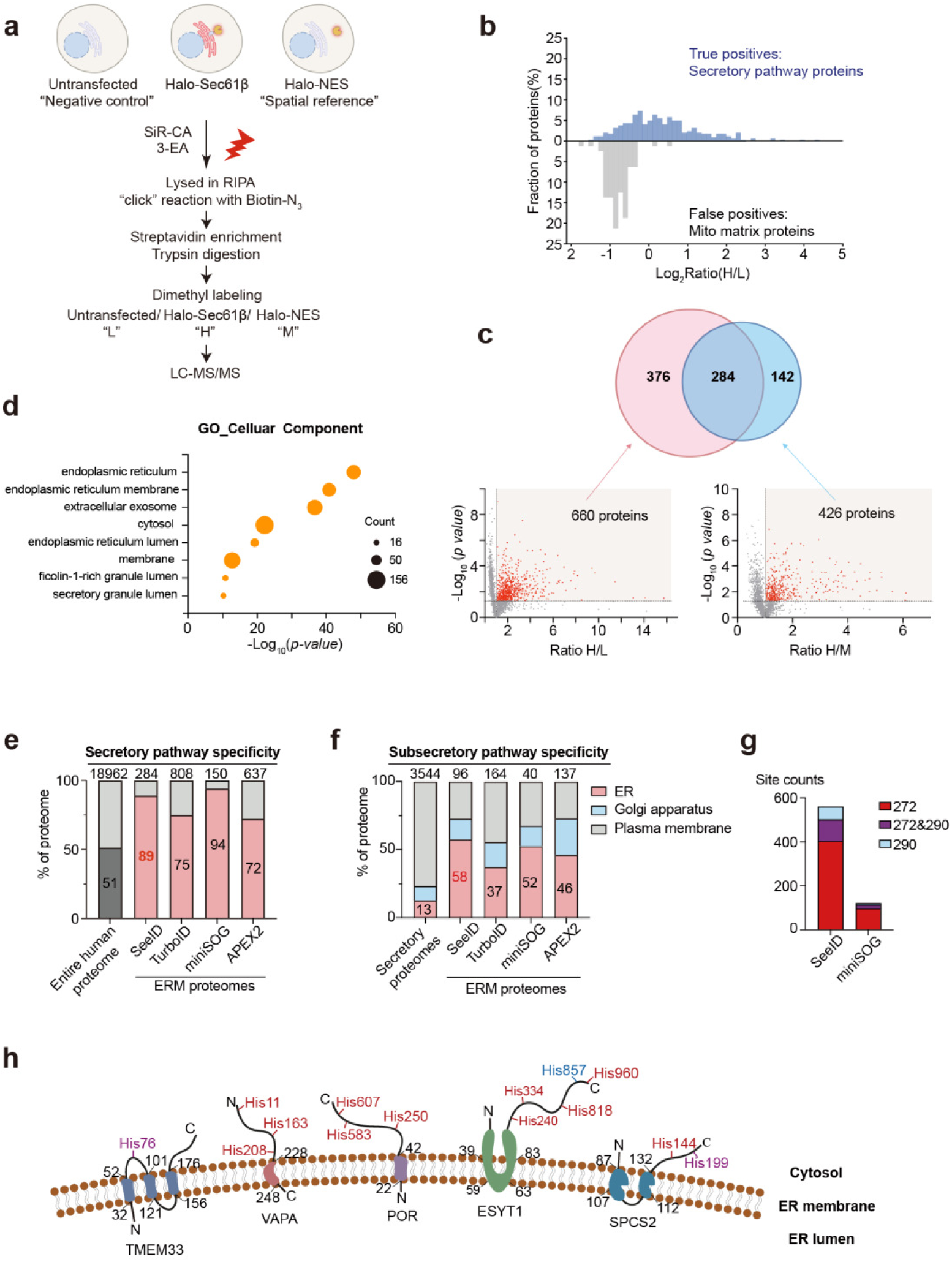
Profiling of the endoplasmic reticulum membrane proteome by SeeID. **a,** Protocol of mass spectrometry-based proteomic analysis of ERM. **b,** Analysis of MS data from the ERM labeling by SeeID. The *Top* histogram displayed the Log_2_Ratio(R/L) distribution of the secretory pathway proteins in the dataset, and the *Bottom* histogram showed the Log_2_Ratio(H/L) distribution of the mito matrix proteins. **c,** Steps for filtering and assigning proteins labeled with SeeID. Enrichment proteins were determined with ratios above 1 and *p-*values below 0.05 in both plots. **d,** GO cellular components analysis of SeeID-enriched proteins. **e-f,** Spatial proteomic comparison of SeeID with other proximity labeling methods at the ERM. **g,** Comparison of the Peptide-Spectrum Matches of histidines labeled by SeeID and miniSOG. **h,** SeeID-labeled sites located at the cytosolic side of ERM. The histidine modification sites are annotated in red.

The SeeID-ERM labeling sample was compared against both the negative control and the spatial reference. True-positive ERM proteins were significantly more enriched than false-positive proteins (e.g., mitochondrial matrix or non-secretory proteins) in both comparisons, demonstrating the high spatial specificity of SeeID (Fig. 3b). SeeID-labeled ERM proteins were defined as those significantly enriched in both comparisons (p < 0.05), yielding a set of 284 high-confidence ERM proteins (Fig. 3c). Gene Ontology (GO) analysis of these proteins revealed significant enrichment for terms related to the endoplasmic reticulum (ER) (Fig. 3d). Additionally, 89% of SeeID-labeled proteins were previously annotated as secretory pathway proteins (Fig. 3e). This specificity was comparable to miniSOG-ERM labeling (94%) and significantly exceeded that of TurboID (75%)^14^ and APEX2^52^ (72%). In a comparative analysis of sub-secretory specificity, SeeID exhibited the highest ER specificity (Fig. 3f). Most labeled ER proteins were confirmed as ERM proteins, with levels comparable to those observed using other PL enzymes (Extended Data Fig. 4b). These results collectively demonstrate that SeeID exhibits excellent spatial specificity.

Previous studies suggest that miniSOG generates singlet oxygen to oxidize histidines in adjacent proteins^20, 47^, a mechanism that is likely shared by SeeID. To investigate this mechanism, the superTOP-ABPP platform was used to identify modification sites of SeeID-ERM and compare them with miniSOG-ERM labeling in parallel (Extended Data Fig. 4c). Streptavidin blotting revealed that SeeID-ERM, activated by NIR light, achieved higher labeling efficiency than miniSOG-ERM under blue light (Extended Data Fig. 4d). Alkyne-labeled peptides were enriched from lysates using agarose beads functionalized with azide groups and acid-cleavable linkers, then identified via LC-MS/MS and analyzed using MSFragger. This approach identified two significant mass shifts on histidines: +272 Da for the 3-EA and 2-oxo-histidine adduct and +290 Da after hydrolysis, observed in both labeling methods (Extended Data Fig.4e). Notably, SeeID labeled more histidines than miniSOG, consistent with the increased labeling observed in streptavidin blots (Fig. 3g and Table S2). Furthermore, an analysis of five ERM proteins with known topologies showed that SeeID labeling sites were predominantly located in regions facing the cytosol (Fig. 3h). Overall, these findings demonstrate that SeeID functions as a genetically encoded photosensitizer with enhanced efficiency and specificity.

We then explored whether SeeID could reveal unknown PPI. The Kirsten rat sarcoma viral oncogene homolog (KRAS) gene is one of the most frequently mutated oncogenes in cancer^53, 54^. Somatic mutations in KRAS results in hyper-activation of downstream MAPK and PI3K-Akt signaling pathways. Despite extensive efforts in studying KRAS and its effector proteins, limited success has been achieved to develop therapeutic strategies targeting KRAS effector proteins ^55–57^. Discovery of new KRAS interacting proteins could provide new insights into KRAS biology as well as potential new strategies targeting KRAS mutant cancers. To test whether SeeID could capture unknown KRAS interacting proteins, we stably expressed Halo-KRAS^WT^ in HeLa cells, and Halo-KRAS^G12C^ in KRAS^G12C^ mutant SW1573 cells. Cells expressing Halo-NES were used as spatial reference control. The correct localization of Halo-KRAS on the plasma membrane was confirmed by confocal microscopy (Extended Data Fig. 5a). Cells expressing the corresponding Halo-KRAS were treated with 1 µM SiR-CA and 500 µM 3-EA, followed by 660 nm LED light irradiation at 50 mW/cm² for 30 minutes. After cell lysis, the biotinylated proteins were then enriched and subjected to LC-MS/MS analysis. To improve peptide coverage, reproducibility and quantitative accuracy, samples were processed with label-free quantitative method and data were acquired using DIA (data-independent acquisition)^58^. 391 and 118 proteins were enriched in HeLa and SW1573 cells respectively, with 82 proteins identified in both cell lines (Fig. 4b,c and Table S3). Of the 427 total enriched KRAS interacting proteins, 215 proteins were reported to associate with KRAS from the BioGRID database, and 212 proteins were newly identified by SeeID (Fig. 4d). According to GO analysis, the enriched proteins were mainly involved in plasma membrane, vesicle and endomembrane organelle, which was consistent with the localization of KRAS (Extended Data Fig. 5b). Reactome pathway analysis showed that these proteins were involved in known signaling pathways related to RAS regulation, including receptor tyrosine kinase signaling, Rho GTPase signaling, MAPK signaling, and transport of small molecules (Extended Data Fig. 5c). Canonical KRAS interacting proteins (e.g. ARAF^59^, RHOA^60^, RICTOR^61^, AFDN^62^ ), as well as proteins identified by previous KRAS proximity labeling efforts (e.g. EGFR, CD44, BSG and ITGB1^61, 63^) were recovered by SeeID, confirming the reliability of our method. For the newly identified 20 proteins both in HeLa and SW1573, we focused on AXL, OSBPL5, and PHLDA1, which were not included in the BioGRID database. AXL has been reported to be involved in the resistance mechanism of anti-EGFR drugs in wild-type RAS patients, and the resistance of KRAS^G12C^-mutant tumor toward KRAS^G12C^ inhibitors^64, 65^. OSBPL5 and other OSBPL family members were found to be up-regulated in Pancreatic ductal adenocarcinoma (PDAC) patient tissue ^66^. PHLDA1 was reported to mediate drug resistance in receptor tyrosine kinase (RTK) driven cancer, and act as an oncogene to promote glioma progression and recurrence^67, 68^ . To further validate the proximity of KRAS to these three proteins, Flag-tagged candidate proteins were co-transfected with HA-tagged KRAS variants in HEK293T cells, followed by proximity ligation assay (PLA) and co-immunoprecipitation. PLA revealed that these proteins produced significant fluorescent signal with either KRAS-WT or KRAS-G12C compared to Flag-NLS control (Fig. 4e). In addition, the interaction of these proteins with KRAS was confirmed by co-immunoprecipitation (Fig. 4f).

**Figure 4.**
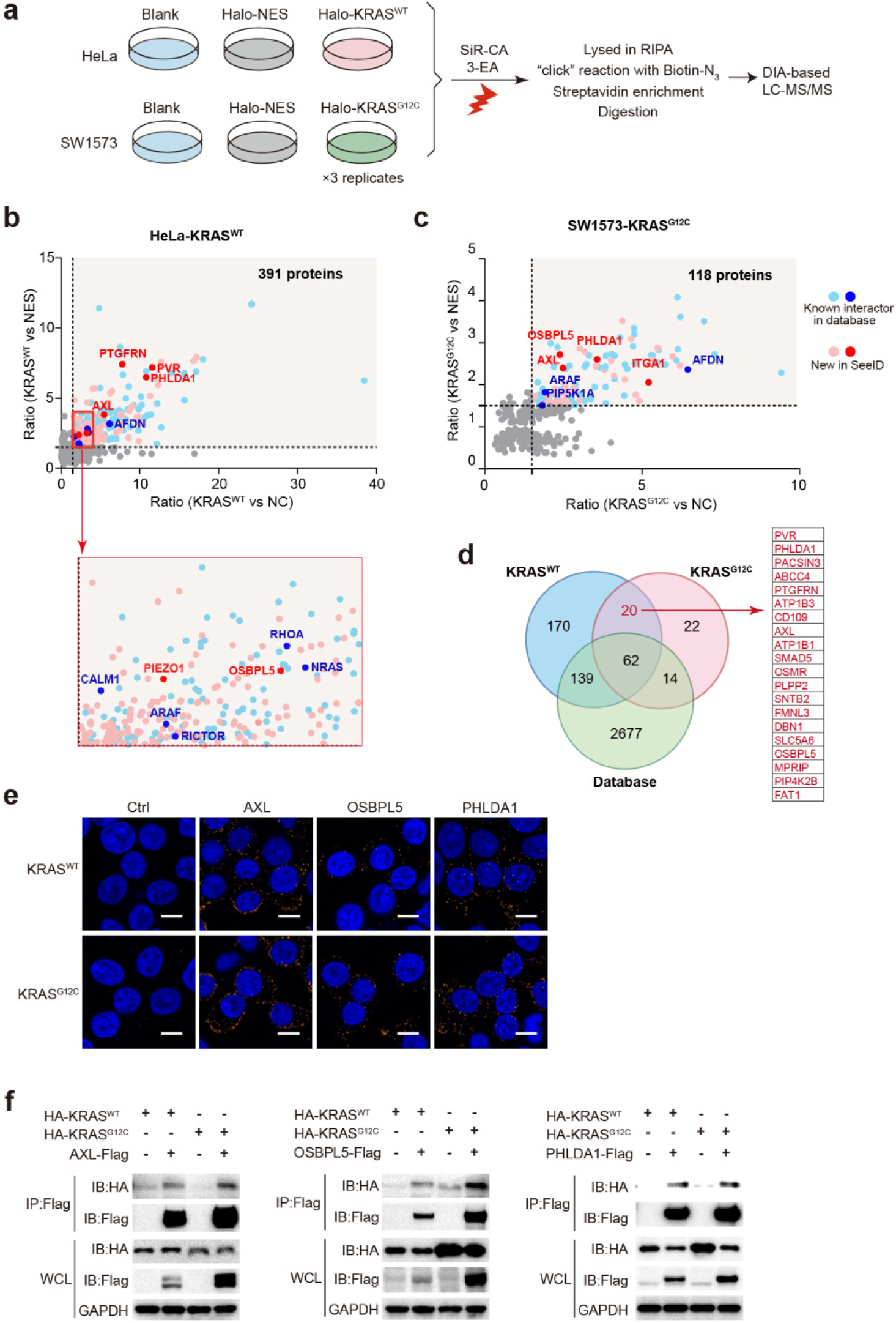
Profiling of KRAS proximity proteome by SeeID. **a,** Protocol of KRAS proximity proteome labeling. **b-c,** Quantitative proteomics volcano plots showed that proteins interacting with wildtype KRAS in HeLa cells (**b**) or G12C mutant in SW1573 cells (**c**). Comparison of Halo-KRAS expression samples with untransfected cells (NC, negative control) or Halo-NES expression cells. Data showed the overlap of proteins with the *p-value*<0.05. 391 and 118 proteins (both including KRAS) were enriched in HeLa or SW1573 cells. **d,** Overlap of identified KRAS interactors by SeeID and BioGRID database. Potential new proteins interacting with both wildtype and mutant KRAS identified by SeeID were listed on the right column. **e,** Proximity Ligation Assay (PLA) in HEK293T cells co-expressing HA-tagged KRAS and Flag-tagged candidate interactors. A Flag-tagged NLS protein was used as a control. Scale bars: 10 μm. **f,** Validation of new KRAS-interacting proteins by co-immunoprecipitation.

Live-cell imaging provides spatial-temporal information on protein dynamics, while PL gives a snapshot of protein interactome at single time point. We thought to develop a method where bulk live-cell imaging events could be analyzed directly by PL. PINK1/Parkin signaling pathway governs mitochondrial quality control via mitophagy^69, 70^. We have previously discovered a small molecule, BL-918, that triggers PINK1 accumulation and Parkin translocation to initiate PINK1/Parkin-mediated mitophagy^71^. To study the BL-918 triggered Parkin translocation process, we treated HeLa cells stably expressing Halo-Parkin with 20 µM BL-918 and tracked the translocation of Parkin by confocal microscopy. Meanwhile, additional dishes with the same cells were prepared for PL. Live-cell imaging showed that Parkin was ubiquitously expressed in the cytosol before BL-918 treatment (Fig. 5a). After 2 hours of BL-918 treatment, Halo-Parkin showed an aggregation pattern, formed vesicle-like structures, and began to translocate to the morphologically altered mitochondria. The correlation became more obvious after 4 hours of drug treatment(Fig. 5a, PCC = 0.44, 0.82, 0.85 at 0, 2, 4 h). In contrast, the localization of Halo-NES was maintained in the cytoplasm after BL-918 treatment (Extended Data Fig. 6a). PL was performed simultaneously at these time points (0 h, 2 h, 4 h) to capture the dynamic proteomic changes of Halo-Parkin as we track the translocation process by live-cell imaging (Fig. 5b). After PL, the samples were first verified by immunostaining then subjected to MS analysis. Consistent with the live cell imaging results, immunofluorescence results of fixed cells after a click reaction with Rhodamine-N_3_ showed that the rhodamine-labeled signal accumulates from being scattered in the cytoplasm to damaged mitochondria, similar to the signal of Halo-SiR (Extended Data Fig. 6b). The interactome of Parkin in this dynamic process was then analyzed by LC-MS/MS using Halo-NES as a spatial control. 269 proteins interacting with Halo-Parkin before BL-918 treatment, and 250 and 93 proteins after BL-918 treatment for 2 h and 4 h, respectively, were obtained (Extended Data Fig. 6c and Table S4). The decreased amount of proteins enriched at 4 hours might result from the lower cell viability at this time point (Extended Data Fig. 6d). Cell viability analysis revealed no significant reduction at 2 hours of BL-918 treatment (Extended Data Fig. 6d). Proteome data revealed a significantly differentiated Parkin interactome upon drug treatment.

**Figure 5.**
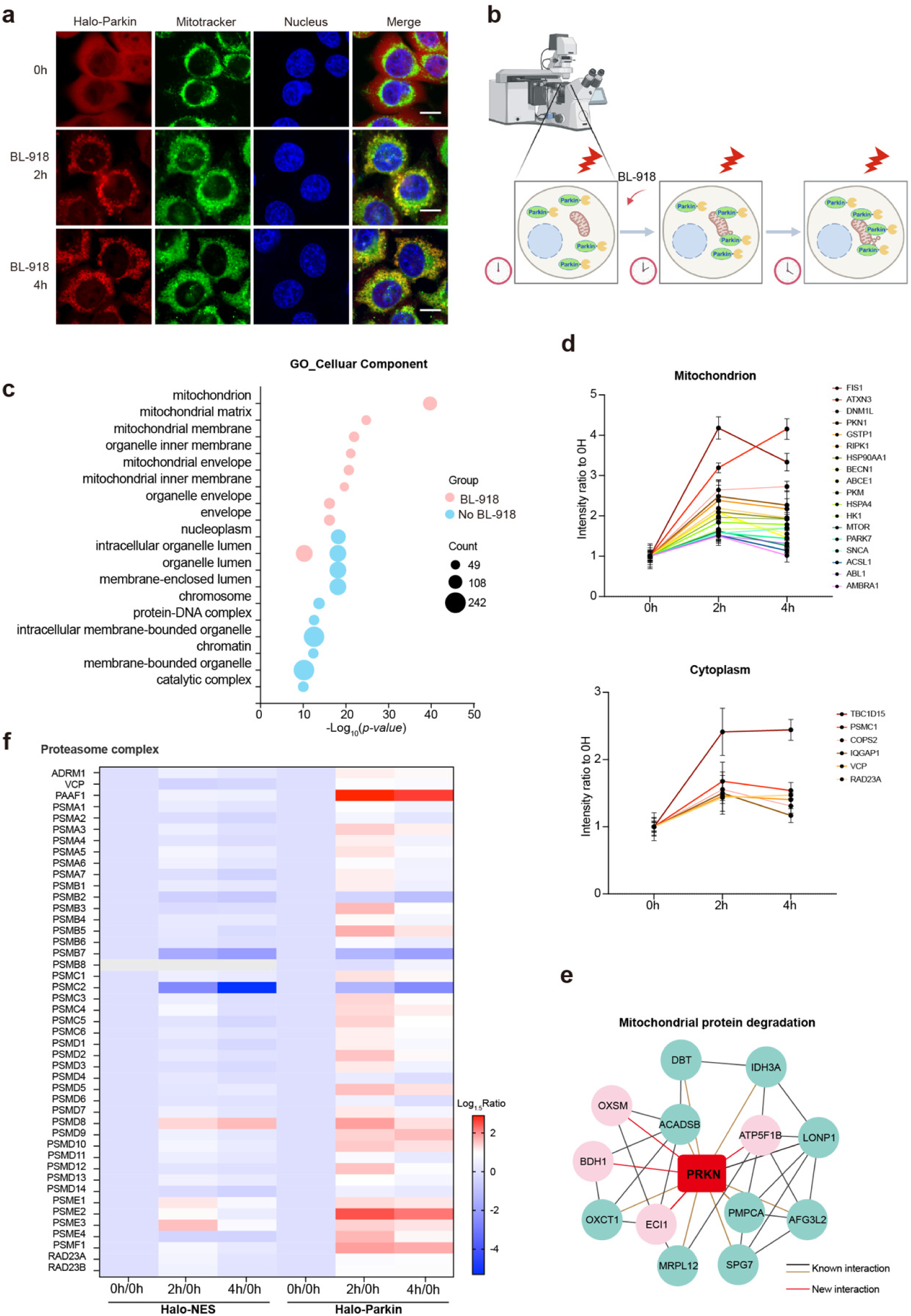
Imaging-guided profiling of Parkin dynamic proteome. **a,** Live cell confocal imaging of HeLa-Halo-Parkin cells treated with BL-918. Cells were treated with 20 μM BL-918 for 0 hour, 2 hours and 4 hours. Mitotracker green were used as Mitochondria marker. Scale bars: 10 μm. **b,** Protocol of imaging-assisted dynamic proteome profiling of Parkin upon treatment with BL-918. **c,** Cellular components in GO enrichment analysis before and after BL-918 treatment. **d,** Interaction network of 13 proteins involving in mitochondria protein degradation pathway with Parkin (Black line: data from STRING, brown line: data from BioGRID, red line: new in SeeID). **e,** The intensity in quantification for representative known Parkin substrates in mitochondrion and cytoplasm. **f,** Heatmap of intensity changes involving in proteasome complex-associated proteins.

Proteins enriched prior to treatment were spread across the nucleoplasm, organelle lumen, nucleus, and other regions, while the mitochondrial membrane was the primary location of proteins enriched after BL-918 treatment (Fig. 5c). The ion intensity of known Parkin interacting proteins was similarly increased with BL-918 treatment. These proteins include PKN1(PINK1), FIS1, HK1, PARK7, MTOR, RIPK1, PKM located in the mitochondria^72–75^, and TBC1D15, PSMC1, VCP, RAD23A in the cytoplasm^72^ (Fig. 5d and Table S5). For the identification of previously unknown interacting proteins of Parkin, we focused on the effectors involved in the mitochondrial protein degradation pathway. 13 proteins in this pathway were identified by SeeID. Among which, 9 of 13 were identified by previous affinity mass spectrometry or proximity labeling^76, 77^, and 4 proteins have not been reported, including ECI1, BDH1, OXSM, ATP5F1B (Fig. 5e). Interestingly, quantitative analysis indicated that 29 out of the 45 proteins associated with the proteasome complex has increased (Ratio >1.5) in ion intensity after 2 hour BL-918 treatment (Fig. 5f). This enrichment was not due to the overall increased protein expression, as the proteasome complex was not enriched in the Halo-NES group. A decrease of proteasome complex enrichment was observed after 4 hours. Although this observation is in line with previous reports indicating that Parkin plays a role in recruiting proteasomes to depolarized mitochondria and facilitating the degradation of ubiquitin-tagged mitochondrial proteins^78, 79^, to the best of our knowledge, the dynamic spatial interplay between the proteasome complex and Parkin during mitophagy has not been discoverd before. This phenomenon demonstrated the dynamic protein interactions during mitophagy, and showcased the potential of imaging-guided PL for studying dynamic protein interactomes.

Application of SeeID in vivo will allow the study protein interactomes in animal models. To demonstrate the utility of SeeID in vivo, we aimed to label the mouse brain. Halo-tagged PDGFR-TM protein (Halo-TM) was used to target SiR to the plasma membrane. Biotin-conjugated probe BA was used as the probe to capture the proximal proteins (Extended Data Fig. 7a). The ability of SeeID to label mouse brain was first demonstrated on acute brain slices. The packaged AAV virus carrying Halo-TM (AAV-Halo-TM) was stereotactically injected to the hippocampus regions of C57BL/6J male mice. After 2 weeks of virus expression in mice, the brains were extracted under anesthesia, submerged in artificial cerebrospinal fluid (ACSF), and cut into 300 µm slices. Brain slices containing hippocampal regions were incubated with 1 µM SiR-CA and 500 µM BA in ACSF for 30 mins on ice, followed by labeling with 660 nm LED light at an intensity of 100 mW/cm^2^ for 45 mins on ice (Fig. 6a). Immunofluorescence demonstrated robust labeling signals in the hippocampus, while the labeling was not observed without Halo-TM expression, SiR-CA or irradiation (Fig. 6b). Western blot results further comfirmed the labeling specificity (Fig. 6c).

**Figure 6.**
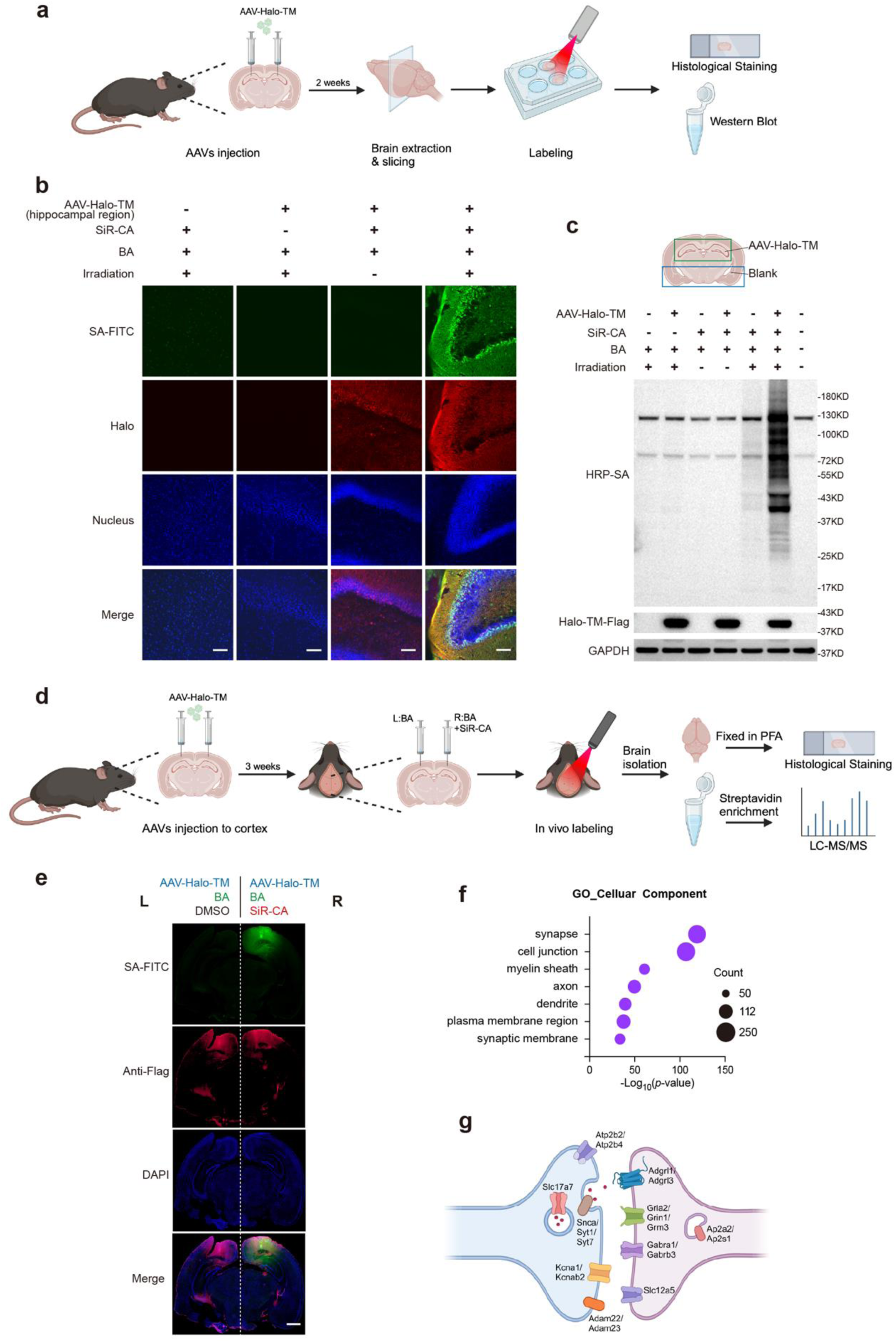
SeeID labeling of mouse hippocampus and cerebral cortex in vivo. **a,** Protocol of proximity labeling in acute brain slices. AAV-Halo-TM was injected into the hippocampus of mice, and the mice were kept for 2 weeks to express Halo-TM in vivo. Mice’s brains were isolated after anesthetization and sliced. The slices were immersed in artificial cerebrospinal fluid and incubated with1 μM SiR-CA and 500 μM BA for 30 minutes, followed by 660 nm LED light irradiation for 45 minutes. **b,** Confocal immunofluorescence images of hippocampus labeling in mouse brain slice. The cortical region was used as non-Halo-TM expressing control. **c,** Western blot analysis of brain slice labeling. The hippocampus and non-hippocampus were dissected and analyzed by streptavidin blotting. **d,** Protocol of in vivo proximity labeling of mouse cerebral cortex. AAV-Halo-TM was injected into the cerebral cortex of mice. After 3 weeks, mice were anesthetized and injected BA alone into the left cerebral cortex or BA and SiR-CA into the right cerebral cortex, followed by 660 nm LED light irradiation for 30 minutes. **e,** Confocal immunofluorescence images of cerebral cortex labeling. The FITC-SA staining indicated the proximity labeling signal and Halo-TM-Flag expression was detected by anti-Flag staining. **f,** GO cellular component analysis of enriched proteins identified in mice cerebral cortex. **g,** Representative hits of the Halo-TM proteome in cerebral cortex synapse.

After confirming SeeID could be utilized for brain slice labeling, we moved to PL in vivo. AAV-Halo-TM was stereotactically injected into the left (L) and right (R) visual cortex region of mice. Labeling of mouse cortical tissues was performed after 3 weeks of in vivo expression. BA or BA and SiR-CA were delivered to the left or right cortical regions of mice at the same site where the virus was introduced by stereotactic injection. Mice were subsequently labelled by 660 nm LED irradiation at 50 mW/cm^2^ for 30 minutes while being anaesthetized (Fig. 6d). Two mice brains after irradiation were dissected after perfusing for subsequent immunofluorescence staining analysis. The cortical tissues of other mice brains were isolated and homogenized to extract proteins for western blot and mass spectrometry analysis. Immunofluorescence results showed a SiR-CA dependent labeling of up to 2 mm tissue depth(Fig. 6e), demonstrating the specificity of our labeling protocol and the advantage of NIR for potential deep tissue labeling. The observation was further confirmed by western blot: the labeling of the right side of the brain tissue of each mouse (with SiR-CA) was significantly higher than that of the left side (without SiR-CA) (Extended Data Fig. 7b).

We next performed mass spectrometry analysis on the above extracted tissue proteins and determined the specificity of in vivo labeling. A total of 515 labeled proteins were enriched when the proteins were filtered in all three biological replicates with Ratio (R/L) >1 (Extended Data Fig. 7c,d and Table S6). GO analysis demonstrated that these proteins were mainly expressed in synapse, cell junction, plasma membrane region, which is consistent with our Halo-TM protein localization (Fig. 6f). The identified proteins were enriched in synaptic structure, e.g. Slc17a7^80^, which regulates presynaptic neurotransmitter transport; Snca^81, 82^, Syt1^83^, Syt7^83^, which control neurotransmitter release; neurotransmitter receptors, e.g. including GABA receptor Gabra1, Gabrb3^84, 85^, Glutamate receptor Gria2^86^, Grin1^87^, Grm3^88^; as well as a number of ion channels, such as Kcna1^89^, Kcnab2^90, 91^ (Fig. 6g). In summary, our experiments demonstrated the utility of SeeID for in vivo PL, and the use of NIR permits spatiotemporally controlled deep tissue labeling, which has not been achieved by PL methods in the literature.

## Discussion

Live-cell fluorescence imaging offers unparalleled spatiotemporal resolution for studying protein dynamics. However, current imaging tools are unable to elucidate molecular mechanisms, particularly the dynamic protein interactomes, underlying imaging observations. In contrast, PL enables the analysis of protein interactomes at a single time point but lacks information about protein dynamics. A tool capable of simultaneously tracking protein dynamics and analyzing protein interactomes would greatly facilitate the study of dynamic protein interactions during biological processes. To address this challenge, SeeID, a method for NIR-activated PL in vitro and in vivo, was developed. SeeID was benchmarked through characterization of various organelle-specific proteomes and the KRAS protein interactome. Leveraging SiR as both a fluorophore and a photosensitizer enabled imaging-assisted analysis of Parkin interactomes upon induction of mitophagy. Considering the ever increasing throughput of proteomics experiments, we envision SeeID would be highly useful for time-lapse analysis of dynamic protein proteomics.

Another major advantage of SeeID is its capability to perform spatiotemporally controlled PL in vivo due to the use of NIR light. Application of current PL methods in vivo is challenging: APEX requires the use of cytotoxic H_2_O_2_, while BioID/TurboID suffers from high background biotinylation. The tyrosinase-based PL strategy shows promise for in vivo studies but is limited by its dependence on copper cations. In addition tyrosinase-based PL do not allow for spatiotemporal control^92^. Blue-light-regulated LOV-Turbo enabled spatiotemporal controlled PL in vivo^19^. PhoxID utilized green light to active photocatalyst 2-monobromofluorescein for PL in vivo^36^. However, the limited tissue penetration of blue/green light restricts their broader utility. Using AAV transduction of Halo-TM in the mouse cerebral cortex, SeeID was shown to label across the cortex (∼2 mm) under 660 nm LED light, demonstrating its potential for deep-tissue PL. In addition, the in vivo labeling specificity was validated by the enrichment of synaptic and plasma membrane proteins.

The cornerstone of SeeID is the discovery that fluorophore SiR could be repurposed as a photosensitizer for ^1^O_2_-mediated oxidation of proximal histidine residues, which would be subsequently captured by an aniline nucleophile for MS analysis. While SiR is widely used as a fluorophore for live-cell super-resolution imaging, its utility as a photocatalyst remains underexplored. Compared to other small-molecule photocatalysts, the fluorogenic nature of SiR enables specific PL of intracellular targets. A recent report demonstrated that the nonfluorogenic photosensitizer Dibromofluorescein nonspecifically labeled cytosolic proteins when used as a PL photocatalyst^48^. Our own data further underscore the importance of fluorogenicity by showing the nonspecific cytosolic labeling of NIR photocatalyst Methylene Blue.

During the preparation of this manuscript, two related light-activated PL methods were published. Engle et al. developed Fluorogen Activating Protein (FAP)-mediated PL of proteins and RNAs^93^. Although this method also utilized NIR light, the specificity of protein PL was not evaluated using MS. Furthermore, the applicability of this system in vivo was not demonstrated. FAP-based fluorescence imaging has been shown to nonspecifically label secretory apparatus, including the nuclear endoplasmic reticulum and the Golgi^94^. ScFv-based FAPs contain internal disulfide linkages and are adapted for use only in nonreducing environments, mainly the cell surface and secretory apparatus. These suboptimal properties of FAP may limit the utility of FAP-based PL methods. Becker et al. reported POCA, a PL method utilizing JF570 as the photocatalyst^48^. While the fluorogenic photosensitizer JF570 enabled PL of the nuclear pore complex, POCA’s utility in vivo was not demonstrated. Moreover, JF570 nonspecifically localized to the mitochondria when conjugated to cholesterol.

Our method is not without limitations. Although clear labeling bands were observed with 660 nm light exposure for 10 minutes (Extended Data Fig. 3c), a photocatalyst with higher ^1^O_2_ generation efficiency could reduce the irradiation time, facilitating the study of dynamic protein proteomics with improved temporal resolution. Efforts to develop more efficient photocatalysts, while retain their fluorescent and fluorogenic properties, are ongoing in our lab.

Overall, we developed SeeID for NIR-activated SiR-enabled PL in vitro and in vivo. Given the widespread utility of HaloTag and SiR fluorophore in the biological community, and the deep tissue penetration ability of NIR light, we envisioned that SeeID is well-positioned for immediate adoption to study dynamic protein proteomics both in vitro and in vivo.

## Methods

### In vitro protein labeling

Halo protein was expressed with an N-terminal hexa-histidine purification sequence. 6xHis-Halo segment was assembled into pET28a(+) by gibson assembly. Protein was expressed in *Escherichia. coli* BL21(DE3) overnight at 16 ℃, extracted by french press at 4 ℃ and were further purified on Ni-NTA agarose (Beyotime) under native conditions, followed by gel filtration on SD-75 using ÄktaPure FPLC instrument (Cytiva). BSA was purchased commercially (Yeasen). For Halo labeling, photocatalyst 10 μM SiR-CA and 100 μM nucleophilic probes were added to a solution of 10 μM Halo in 100 μL PBS buffer. For BSA labeling, 10 μM SiR-CA or MB and 100 μM nucleophilic probes were added to a solution of 5 μM BSA with or without 10 μM Halo in 100 μL PBS buffer. The mixtures were added into 1.5 mL colorless EP tubes and irradiated with 200 mW/cm^2^ 660 nm LED light at 4 ℃ for the indicated time. After the biotin-conjugated probes labeling, 40 μL samples were removed and combined with 10 μL 5×SDS loading buffer for subsequent western blot detection. After the alkynyl probes labeling, samples were washed twice with PBS in Ultra Centrifugal Filter (10 kDa MWCO) and recovered with 100 μL PBS. 50 μL recovered proteins were mixed with 100 μM Biotin-N_3_ in the presence of 1 mM BTTAA, 500 μM CuSO_4_, 2.5 mM sodium ascorbate, and subjected to click reactions on Thermo Shaker at 25 ℃, 1000 rpm. Samples were washed in Ultra Centrifugal Filter again and recovered with 50 μL PBS, adding 5×SDS loading buffer for western blot detection.

### Singlet oxygen detection

Singlet oxygen generation by SiR-CA upon 660 nm light irradiation was detected by the commercially available Singlet Oxygen Sensor Green (SOSG) (Beyotime). 1 μM SOSG probe was added to the in vitro protein labeling sample. After irradiation, 100 μL solution per sample was transferred to a 96-well black plate immediately and the fluorescence signal was measured by TECAN-Spark microplate reader. To quench the singlet oxygen, 5 mM sodium ascorbate was added to the sample before irradiation.

### Cell culture and plasmid transfection

HEK293T, HeLa, U2OS, and SW1573 cells were cultured in Dulbecco’s modified Eagle’s medium (DMEM, Gibco) supplemented with 10% FBS and 1% Pen-Strep at 37 ℃, 5% CO_2_. Plasmids with segments of Flag-Halo-NLS, Flag-Halo-NES, Flag-Halo-Sec61β, TOMM20-Halo-Flag, LAMP1-Halo-Flag, and Flag-miniSOG-Sec61β were cloned into pCDNA3.1 expression vector by Gibson assembly for transient transfection expression. Transfection for transient protein expression was performed using the NEOFECT™ DNA transfection reagent (Neofect) in HeLa and U2OS, or Lipo8000™ Transfection Reagent (Beyotime) in HEK293T according to the manufacturer’s instructions.

### Construction of stable cell lines

Segments of Flag-Halo-NES, Flag-Halo-KRAS, Flag-Halo-KRAS(G12C), and Flag-Halo-Parkin were inserted into pDest9 vector. The constructed pDest9 plasmids were co-transfected with psPAX2 and pMD2.G into HEK293T using Lipo8000™ Transfection Reagent. Viral supernatant was collected 48 hours after transfection and transduced to the indicated cell line for 48 hours. Cells were selected using the complete culture medium with 2.5 μg/mL puromycin after transduction for approximately 1 week, subsequently maintained in the complete culture medium with 1 μg/mL puromycin.

### Cell viability assay

Cells that were blank or transfected with plasmids for 8 hours, were digested and inoculated 5000 cells per well on a 384-well white plate. Cell viability was detected after being treated with different concentrations of photocatalyst and probe, or different irradiation times and intensities, using Cell Counting-Lite 2.0 (Vazyme). The luminescent signals were acquired by TECAN-Spark microplate reader.

### Labeling in living cells

For SeeID labeling, HEK293T and HeLa cells were cultured in 6 well plates following the transfection of Halo-fusion protein for 24 hours and washed twice with PBS. Cells were treated with SiR-CA (or other catalyst) diluted in HBSS buffer for 1 hour at 37 ℃, washed with PBS twice, and incubated with 3-EA (or other probe) in HBSS buffer for 10 minutes. Irradiated the cells under 660 nm LED of 50 mW/cm^2^ intensity at room temperature for 30 minutes. The labeled cells were washed with PBS twice and lysed in EDTA-free RIPA lysis buffer (50 mM Tris-HCl pH 8, 150 mM NaCl, 0.1% SDS, 0.5% sodium deoxycholate, 1% Triton X-100, 1× protease inhibitor cocktail) on ice for 30 minutes. Samples were centrifuged at 20,000 g for 10 minutes at 4 ℃ and the protein concentration was normalized using BCA Protein Assay Kit (Yeasen). For biotin-conjugated probe labeling, the adjusted lysates were added with 5×SDS loading buffer directly followed by western blot analysis. For alkynyl probe labeling, the clarified supernatants were subjected to click reaction with 100 μM Biotin-N_3_ for immunoblotting analysis or 100 μM N_3_-Rhodamine or in-gel fluorescence, in the presence of 1 mM BTTAA, 500 μM CuSO_4_ and 2.5 mM sodium ascorbate at 25 ℃ for 2 hours. After the click reaction, the proteins were extracted using chloroform-methanol precipitation and dissolved with PBS containing 0.5% SDS.

For miniSOG labeling, HEK293T cells were transfected with Flag-miniSOG-Sec61β for 24 hours, followed by incubation with 500 μM 3-EA in HBSS for 10 minutes, and irradiated under 450 nm blue LED for 20 minutes. The labeled cells were subsequently treated in the same manner as the process of SeeID labeling.

### In-gel fluorescence and western blot

The protein samples were mixed with 5×SDS loading buffer and boiled at 95 ℃ for 10 minutes. Samples were loaded into 4 to 20% Hepes-Tris 15-well precast gels, and electrophoresis was run in 1× Precast Running Buffer (Yeasen) at a constant 100-120 volts for 60∼90 minutes.

For in-gel fluorescence, gels were transferred to water and scanned by Tanon 5200multi. Coomassie Brilliant Blue (CBB) staining served as the loading control.

For western blot, the protein within gel was transferred to 0.45 μm-PVDF membrane (Millipore) at a constant 300mA current for 1 hour. Membranes were blocked with 5% milk diluted in 1×TBST buffer for 2 hours at room temperature with gentle rocking. HRP-conjugated streptavidin (Beyotime) or HRP-conjugated Flag (LabLead) was diluted in TBST with 5% milk and incubated with membranes for 2 hours at room temperature. The membranes were washed 3 times in TBST and exposed to Hith-sig ECL Western Blotting Substrate (Tanon). The imaging data were collected using Tanon 4600 Gel Image System. Primary antibodies were diluted in Antibody Dilution Buffer (Beyotime) and incubated with membranes overnight at 4 ℃. The membranes were washed 3 times and incubated with secondary antibodies diluted in TBST for 1 hour at room temperature. The imaging data were acquired after another 3 washes.

Primary antibodies used in this study showed as following: Anti-Flag (Abclonal, AE005), Anti-HA (Abclonal, AE008), GAPDH (Proteintech, 60004-1-1g).

### Immunofluorescence

HeLa, U2OS or HEK293T was inoculated on 35mm glass coverslips (Cellvis) 16 hours before transfection with the corresponding plasmid. 24 hours after transfection, cells were treated with 1 μM SiR-CA ( or other catalysts) in HBSS for 1 hour, washed twice with PBS, and incubated with 500 μM 3-EA (or other probes) in HBSS for 10 minutes. Cells were irradiated under 50 mW/cm^2^ 660 nm LED for 20 minutes, washed twice with PBS, and fixed with pre-cold methanol (-20 ℃) at -20 ℃ for 20 minutes. After another 3 washes with PBS, 200 μL click reaction buffer (100 μM Biotin- N_3_ or Rhodamine-N_3_, 1 mM BTTAA, 500 μM CuSO_4_, and 2.5 mM sodium ascorbate) was added to the area of central circle and incubated at room temperature for 1 hour. Cells were washed 3 times with PBS and blocked with 5% BSA in PBS for 1 hour at room temperature. Cells clicked with Biotin-N_3_ were stained with FITC-Streptavidin (APExBIO) at a dilution of 1:500 in 5% BSA at room temperature for 1 hour, followed by 3 washes with PBST, and treated with DAPI (1 μg/mL) in PBST for 10 minutes to nucleus staining. Cells clicked with N_3_-Rhodamine were stained with DAPI directly. The cells were maintained in PBS for subsequent imaging by Olympus FV-1200.

### Live cell imaging

Cells were inoculated into 35mm glass bottom dish 1 day before treatment. HeLa stably expressing Halo-KRAS^WT^ and SW1573 stably expressing Halo-KRAS^G12C^ were treated with 1 μM SiR-CA in HBSS for 1 hour and washed twice with PBS before imaging. HeLa stably expressing Halo-NES and Halo-Parkin were continuously treated with 20 μM BL-918 for 0, 2, or 4 hours, 1 μM SiR-CA along with 500 nM Mito-Tracker Green (Beyotime) and Hoechst 33342 (Yeasen) were added at 1 hour before finish time. Images were captured by Olympus FV-1200 with live cell workstation.

### Proximity Ligation Assay

HEK293T cells were inoculated into 4-chamber glass bottom dish. The HA-tagged KRAS^WT^/ KRAS^G12C^ plasmid was co-transfected with Flag-tagged candidate interactors, using a Flag-tagged NLS plasmid as a control. 24 hours after transfection, cells were washed with PBS once prior to fixation with pre-cold methanol (-20 ℃) at -20 ℃ for 20 minutes. Duolink® In Situ Orange Starter Kit Mouse/Rabbit (Millipore Sigma) was used according to the manufacturer protocol for PLA.

### Co-Immunoprecipitation

HEK293T cells were cultured in a 6-well plate and allowed to reach ∼60-70% confluency before co-transfected with HA-tagged KRAS and Flag-tagged candidate proximal proteins. The cells were lysed in 200 μL RIPA 24 hours after transfection, and the cell lysates were incubated at 4 ℃ for 30 minutes before clarification by centrifugation at 15,000 g for 10 minutes at 4 ℃. Anti-Flag magnetic beads (LabLead) were rinsed twice in RIPA using 30 μL of beads for each harvested well. A 20 μL aliquot of the cell lysates was collected as input, and the remaining supernatant was added RIPA up to 500 μL and incubated with the beads overnight at 4 ℃ with rotation. The beads were washed three times with RIPA for 10 minutes , and the immunoprecipitants were resuspended in 2×SDS loading buffer. Protein was eluted by heating at 95 °C for 10 min before analysis of protein content by immunoblotting.

### Samples preparation for mass spectrometry

For ERM proteome analysis, HEK293T cells were inoculated into 15 cm dishes one day before transfection. Three dishes per group were used as biological replicates. The following day, the cells were either left untransfected or transfected with Flag-Halo-NES and Flag-Halo-Sec61β, respectively. 24 hours after transfection, cells were washed once with PBS and treated with 1 μM SiR-CA diluted in HBSS for 1 hour at 37 °C. Next, cells were washed twice with PBS and treated with 500 μM 3-EA in HBSS for 10 minutes at 37 °C. Subsequently, the cells were irradiated under 660 nm LED light for 30 minutes at room temperature.

For KRAS interactome analysis, blank of HeLa, stable cellines of HeLa-Halo-NES and HeLa-Halo- KRAS^WT^, blank of SW1573, SW1573-Halo-NES and SW1573-Halo-KRAS^G12C^ were plated into 15 cm dishes one day before labeling. Cells were labeled as described above. For Parkin dynamic proteome analysis, blank of HeLa and stable cellines of HeLa-Halo-NES, HeLa-Halo-Parkin were plated into 15 cm dishes one day before drug treatment. Cells were treated with 20 μM BL-918 in HBSS for 0, 1, 3 hours, followed by addition of 1 μM SiR-CA for additional 1 hour. Then the cells were washed twice with PBS to remove the redundant SiR-CA, and treated with 500 μM 3-EA in HBSS for 10 minutes at 37 °C. Cells were also labeled as described above.

After irradiation, the labeled cells were washed twice with PBS, scraped off with a cell scraper, and centrifuged in 15 mL centrifuge tube at 4 °C. Then the cells were lysed in 2 mL EDTA-free RIPA lysis buffer on ice for 30 min. Samples were centrifuged at 20,000 g for 10 mins at 4 °C and the protein concentration was normalized to a final protein concentration of 2 mg/mL using BCA Protein Assay Kit. 500μL lysate was removed and subjected to click reaction with Biotin-N_3_ via CuAAC for 2 h at room temperature.

After the click reaction, the proteins were extracted using chloroform-methanol precipitation and dissolved in RIPA buffer. 50 μL streptavidin magnetic beads were washed with RIPA once and co-incubated with samples on a rotary shaker overnight at 4 ℃. Pellet the beads on a magnetic rack and wash the beads twice with 1 mL RIPA buffer, once with 1 mL of 1 M KCl, once with 1 mL of 0.1 M Na_2_CO_3_, once with 1 mL of 2 M urea in 10 mM Tris-HCl (pH 8.0), and twice with RIPA buffer. The enriched proteins with magnetic beads were washed twice by 1 mL PBS, the resulting beads were resuspended in 500 μL 100 mM triethylammonium bicarbonate (TEAB) buffer with 6 M urea and 10 mM dithiothreitol (DTT) at 35 °C for 30 minutes and alkylated by addition of 20 mM iodoacetamide (IAA) at 37 °C for 30 minutes in the dark. The beads were washed with 1 mL of 100 mM TEAB buffer, and resuspended in 200 μL of 100 mM TEAB buffer with 2 M urea/PBS, 1 mM CaCl_2_ and 10 ng/μL trypsin. Trypsin digestion was performed at 37 °C on a thermomixer overnight. Collect the digested peptides in fresh microcentrifuge tubes, wash the the beads twice with 50 μL of 100 mM TEAB buffer and the supernatant was combined with the peptide solution.

For dimethyl labeling-based quantitative proteomics of ERM proteome, the peptides were reacted with 12 μL of 4% D^13^CDO , DCDO, or HCHO for “heavy”, “medium” or “light” labeling respectively. 12 μL of 0.6 M NaBH_3_CN was added in “light” or “medium” labeled samples, 12 μL of 0.6 M NaBD_3_CN was added in “heavy” labeled samples. The solutions were reacted for 2 hours at room temperature and terminated by 48 μL of 1% ammonium hydroxide for 15 minutes. 24 μL of 2% formic acid (FA) was added to neutralize the remaining ammonium hydroxide. The peptides were fractionated by suspension and desalted using homemade C18 tips. The peptides were then sequentially fractionated using 6%, 12%, 15%, 18%, 21%, 25%, 30%, 35%, 50%, 80% and 100% acetonitrile (ACN) with 10 mM ammonium hydrogen carbonate (ABC) (pH=10). The eluate containing the peptides was collected for LC-MS/MS analysis.

For investigation of KRAS interactome and Parkin dynamic proteome, the eluted peptides were desalted on C18 StageTips: The desalting column was activated and equilibrated by CAN and 1% FA water respectively. The peptide samples were loaded onto the column three times. Then, 1 mL of 1% FA in water was added and this washing step was repeated three times. Subsequently, 300 μL of 30%, 50%, and 80% ACN were added, and the eluted peptides were combined.

For in vivo labeling samples, the proteins were enriched on-tip. Briefly, the homemade C18 tips were blocked with 60 μL 2% SDS and centrifuged at 3000 g for 3 minutes. The sepharose beads slurry, prewashed twice with RIPA lysis buffer, were loaded onto the tips and centrifuged at 1,000 g for 3 minutes. About 100 μg sample was then loaded onto the tip and centrifuged at 100 g for 1 hour. After biotinylated proteins were captured, the tips were washed 4 times with 60 uL of RIPA buffer at 1,500 g for 1 minute. Then C18 was activated with 60 μL of 0.5 % (v/v) acetic acid (HOAc) in 80 % ACN by centrifugation at 6000 g for 3 mins. For reduction of cysteine residues, 20 μL of 50 mM ABC containing 10 mM DTT was added and incubated at 25°C with 600 rpm for 15 minutes, and removed with centrifugation at 6,000 g for 3 minutes. 10 μL digestion buffer containing 0.25 μg/μL trypsin, 50 mM lAA, and 50 mM ABC was loaded onto the tips and the samples were incubated at 37°C for 1 hour with 600rpm in the dark to digest and alkylate the proteins. The digested peptides (retained on C18) were washed with 60 μL 10 mM NH_4_HCO_3_ three times. Afterward, the peptides were eluted with 60 μL of 0.5 % (v/v) HOAc in 80 % ACN by centrifugation at 5,000 g or 5 minutes three times.

### SuperTOP-ABPP

The same cell lysate samples from ERM proteome analysis were collected. The proteins were precipitated with chloroform-methanol, and the protein pallets were resuspended in 0.4% SDS PBS to a final protein concentration of 2 mg/mL. The proteins were then reduced with 10 mM DTT at 37 °C for 30 minutes and alkylated with 20 mM IAA at 35 °C for 30 mins in dark. The alkylated proteins were precipitated with chloroform-methanol again and then resuspended in PBS buffer with ultrasonication. The protein was digested with trypsin for 16 h at 37 °C and mixed with acid-cleavable azide-functionalized beads in the presence of 2 mM BTTAA, 1 mM CuSO_4_, and 0.5% sodium ascorbate at 29 °C for 2 hours. After the click reaction, the beads were washed once with 1 mL of 8 M urea, twice with 1 mL of PBS and 5 times with 1 mL of H_2_O. Peptides were cleaved on a thermomixer using 200 μL of 2% FA for 30 minutes twice. Centrifuge at 2000 g for 5 minutes to collect the peptide eluate for LC-MS/MS analysis.

### Animals

Male C57BL/6J mice at 7-8 weeks of age were obtained from Beijing Vital River Laboratory Animal Technology Co., Ltd (Beijing, China) and gifted by the Laboratory Animal Center of the Institute of Genetics and Developmental Biology, Chinese Academy of Sciences (Beijing, China). Mice were housed in pressurized, individually ventilated cages (PIV/IVC) and maintained under specific-pathogen-free conditions, with free access to food and water in a 12 h light/dark cycle. All animal studies were approved by the the Institutional Animal Care and Use Committees of Tsinghua University (Beijing, China) and Animal Care and Use Committee of Institute of Genetics and Developmental Biology, Chinese Academy of Sciences (Beijing, China).

### Mice intracranial surgery

Mice were anesthetized with isoflurane inhalation, placed in a stereotaxic frame (RWD life science), and then were injected 300 nL/lateral or 1000 nL/lateral AAV virus (AAV-Halo-TM, 3.47x10^12^ vg/mL) into the visual cortex and the hippocampus respectively, using the glass microelectrode needle (RWD life science). Stereotactic coordinates used for the visual cortex were 3.08 mm posterior to the bregma (A.P: -3.08mm), 1.5 mm lateral to the midline (ML:±1.5 mm) and 0.5 mm deep (DV: -0.5mm), and for the hippocampus were 1.8 mm posterior to the bregma (A.P:-1.8 mm), 1.25 mm lateral to the midline (ML:±1.25 mm) and 1.55-1.8 mm deep (DV: -1.55 mm and -1.8 mm). After infusion of the regents, the syringe needle was kept in place for 10 mins to minimize the backflow of the regents. After 2-3 weeks of virus expression in mice brains, labeling experiments were performed.

For acute brain slices labeling, mice infected with the AAV in the hippocampus were anesthetized with Avertin (500 mg/kg, i.p. injection, Sigma), and their brains were quickly dissected out and placed in ice-cold carbogenated (5% CO_2_, 95% O_2_) artificial cerebrospinal fluid (ACSF) containing (mM): choline chloride (110), KCl (2.5), NaH_2_PO_4_ (1.25), myo-inositol (3), sodium pyruvate (3), NaHCO_3_ (25), MgCl_2_ (3), CaCl_2_ (0.1). 300 μm coronal slices were cut on a Leica vibratome (Leica, VT1200S). 6-8 slices per group were plated in 12-well plate soaking in ACSF addition with SiR-CA (1 μM) and BA (500 μM) under oxygenated conditions on ice. The slices were irradiated under 660 nm LED of 200 mW/cm^2^ intensity for 45 minutes. After irradiation, the slices were washed 3 times with PBS for 10 mins each. For protein extraction, the hippocampal and non-hippocampal tissues were dissected from the slices and processed in PBS with tissue homogenizer. The samples were lysed by adding RIPA lysis buffer at 4 °C for 30 minutes. Lysates were centrifuged at 20,000g for 10 min at 4°C, following with BCA quantification and western blot analysis. For histology, the brain slices were washed 3 times in PBS following overnight fixation in 4% paraformaldehyde (PFA), then incubated in 10% normal donkey serum (NDS) in PBS with 0.3% Triton X-100 (PBST) for 4 hours at room temperature. The slices were stained in 5% NDS-PBST with FITC-Streptavidin (1:500) at 4 °C for two overnights. Samples were washed 3 times in PBST at room temperature and incubated in DAPI (1:1000) in PBST for 30 minutes, subsequently washed twice in PBST and once in PBS. The slices were mounted on glass slides in Fluoromount G (Yeasen), followed with glass coverslips mounting on slides and sealing the edges with clear nail polish. The slides were stored at 4 °C overnight, images were acquired by Olympus FV1200.

For labeling *in vivo*, mice infected with the AAV in the visual cortex were subjected to intracranial surgery again. For the injection, 300 nL PBS with BA (10 mM), or 300 nL PBS with BA (10 mM) and SiR-CA (200 μM), was administered into the left and right visual cortex, respectively. Mice were maintained anesthetic, and irradiated with 660 nm LED light above the head for 30 minutes. Saline was administered to the exposed skull at 10-minute intervals to keep it from drying. For protein extraction, the mice brain was dissected out after euthanasia. The right and left visual cortical regions of the labeled site were isolated respectively and filled into fresh EP tubes. Samples were homogenized and lysed RIPA buffer, followed by centrifugation. A small amount of sample was taken for western blot assay, the rest lysates were stored at -80 °C. For histology, the mice were perfused with PBS and 4% PFA in sequence. Brains were dissected and post-fixed in 4% PFA overnight at 4°C, and then transferred to 30% sucrose in PBS overnight at 4 °C. Dehydrated brain tissues were then embedded in O.C.T. compound (Tissue-Tek) and frozen on dry ice. Serial sections were coronally prepared at 14 μm for cryostat sections. For IF staining on tissues, frozen brain sections were oven dried at 42 ℃ for 30 mins, The following immunofluorescence procedures were the same as acute brain slices. Images were acquired by Olympus VS200.

### LC-MS/MS

Peptides were separated using a loading column (100 µm × 2 cm) and a C18 separating capillary column (100 µm × 15 cm) packed in-house with Luna 3 μm C18(2) bulk packing material (Phenomenex, USA). The mobile phases (A: water with 0.1% formic acid and B: 94% acetonitrile with 0.1% formic acid) were driven and controlled by a Vanquish™ Neo UHPLC system (Thermo Fisher Scientific). The LC gradient for protein samples was held at 4% for the first 1.8 min of the analysis, followed by an increase from 4.5 % to 5% B from 1.8 to 2 min, an increase from 5% to 20% B from 2 to 54 min, an increase from 20% to 35% B from 54 to 78 min and an increase from 35% to 99% B from 78 to 81.5 min. The LC gradient for superTOP-ABPP samples was held at 4% for the first 4 min of the analysis, followed by an increase from 5 % to 20% B from 4 to 109 min, an increase from 20% to 35% B from 109 to 150 min and an increase from 35% to 99% B from 150 to 159 min.

For the samples analyzed by Q Exactive-plus series Orbitrap mass spectrometers (Thermo Fisher Scientific), the precursors were ionized using an EASY-Spray ionization source (Thermo Fisher Scientific) source held at +2.0 kV compared to ground, and the inlet capillary temperature was held at 320 °C. Survey scans of peptide precursors were collected in the Orbitrap from 350-1800 Th with an AGC target of 3,000,000, a maximum injection time of 20 ms and a resolution of 70,000. For dimethyl labeling and SuperTOP-ABPP samples, the data-dependent acquisition mode was selected. Monoisotopic precursor selection was enabled for peptide isotopic distributions, precursors of z = 2–7 were selected for data-dependent MS/MS scans, and dynamic exclusion was set to 25 s with a ± 10 ppm window set around the precursor monoisotope. For KRAS, Parkin and in vivo samples, the data-independent acquisition mode was selected. For each DIA window, resolution was set to 17,500. AGC target value for fragment spectra was set at 1,000,000 with an auto IT. Normalized CE was set at 28%. Default charge was 3 and the fixed first mass was set to 200 Th.

### Data analysis

For dimethyl labeling data analysis, the MS/MS data were processed with the MaxQuant software (v2.4.11.0) with the Human UniProt isoform sequence database (3AUP000005640). Carbamidomethyl cysteine was set as fixed modification, and methionine oxidation and acetyl N-terminal were set as variable modifications. DimethLys0 and DimethNter0 were enabled in the light labels, DimethLys4 and DimethNter4 were enabled in the medium labels. DimethLys8 and DimethNter8 were enabled in the heavy labels. The minimal peptide length was set to 7 amino acids, and the maximal mass tolerance was set to 20 ppm and 4.5 ppm in the first search and main search. The false discovery rate (FDR) was set to 1%. Proteins with at least two unique peptides were reported as identified proteins. For protein quantification, the functions of “re-quantify” and “match between runs” were enabled, and proteins with at least two quantified peptides (at least one unique peptide) were defined as quantified proteins. After the program finished, dimethyl labeling data analysis depended on the output file “combined-txt- proteinGroups.txt”, including Ratio H/L, Ratio H/M, and so on. Based on the above information, we can sort out the dimethyl labeling quantification and obtain the corresponding protein information.

For SuperTOP-ABPP data analysis, the MS/MS data were processed with the MaxQuant software (v2.4.11.0) with the Human UniProt isoform sequence database (3AUP000005640). Methionine oxidation, protein N-terminal acetylation, and probe modification on histidine were set as variable modifications, while carbamidomethylation of cysteine was set as a static modification. “Trypsin” was selected as the digestion enzyme with a maximum of 2 missed cleavages. All other parameters were set as default. Modified sites with localization probability ≥ 0.75 were selected for further analysis.

For DIA data analysis, the raw data were processed using DIA-NN in an advanced library-free module. The main search settings for in silico library generation were set as following: trypsin/P with maximum 3 missed cleavage; protein N-terminal M excision on; carbamidomethyl on C as fixed modification; oxidation on M as variable modification; peptide length from 7-30; precursor charge 1-4; precursor m/z from 300 to 1800; fragment m/z from 200 to 1800. The Human UniProt isoform sequence database (3AUP000005640) was used to annotate proteins for human cell samples, and the Mouse UniProt isoform sequence database (3AUP000000589) was used to annotate proteins for mouse samples. Other search parameters were set as following: quantification strategy was set to ‘‘QuantUMS (high precision)’’ mode; cross-run normalization was off; MS2 and MS1 mass accuracies were set to 0, allowing the DIA-NN to automatically determine mass tolerances; Scan window was set to 0 corresponding to the approximate average number of data points per peak; Peptidoforms and MBR were turned on; neural network classifier was single-pass mode.

### Statistical analysis

Gene ontology analysis was conducted using the DAVID and g:Profiler databases, for the ERM proteome and other proteome data, respectively. The Reactome pathway for KRAS interactors and the Parkin network were mapped using the Cytoscape software, version 3.10.3. The heatmap, bar and line charts in the figures were generated using GraphPad Prism 10. Pearson’s correlation coefficient was analyzed using the ImageJ software. The schematic workflows were created in https://BioRender.com. Significance was defined as a \**p* < 0.05, \*\**p* < 0.01 and \*\*\**p* < 0.001 for the two-tailed *Student’s t*-test. At least two independently replicates were performed for all experiments with similar results.

## Acknowledgements

Financial support for this work was provided by National Key Research and Development Program of China (No. 2021YFA0910900, L.C.; No. 2024YFA1308000, W.Q.), National Natural Science Foundation of China Science (22477066 and 92478128, W.Q.; 32300614, X.W.) “Thousand Youth Talents Plan” (L.C.), the Fundamental Research Funds from Beijing National Laboratory for Molecular Sciences (BNLMS202301, W.Q.), the Shenzhen Medical Research Fund (B2401004, W.Q.), Beijing Frontier Research Center for Biological Structure (L.C.; W.Q.), startup funding and “Dushi Plan” from Tsinghua University (L.C.; W.Q.), Center for Life Sciences postdoctoral fellowship (W.W.; X.P.). We thank Yanli Zhang (THU) for helping with confocal imaging experiments. We thank Prof. Guotai Xu (NIBS) for sharing of the SW1573 cell line.

## Author Contributions Statement

L.C. and W.Q. conceived the project. W.W. developed the PL protocol, performed the PL in cellulo and in vivo, and analyzed the MS data. H.G. performed the MS experiments, processed and analyzed the MS data, with the help from X.P.. X.Y. synthesized and characterized the chemical probes, and performed the PFAA reduction reaction. X.W., Y.R., Z.B., L.Z., Z.W. and X.Z. helped with the in vivo experiments. L.C., W.Q. and W.W. wrote the manuscript with feedbacks from all other authors.

---

**Extended Data Fig. 1.**
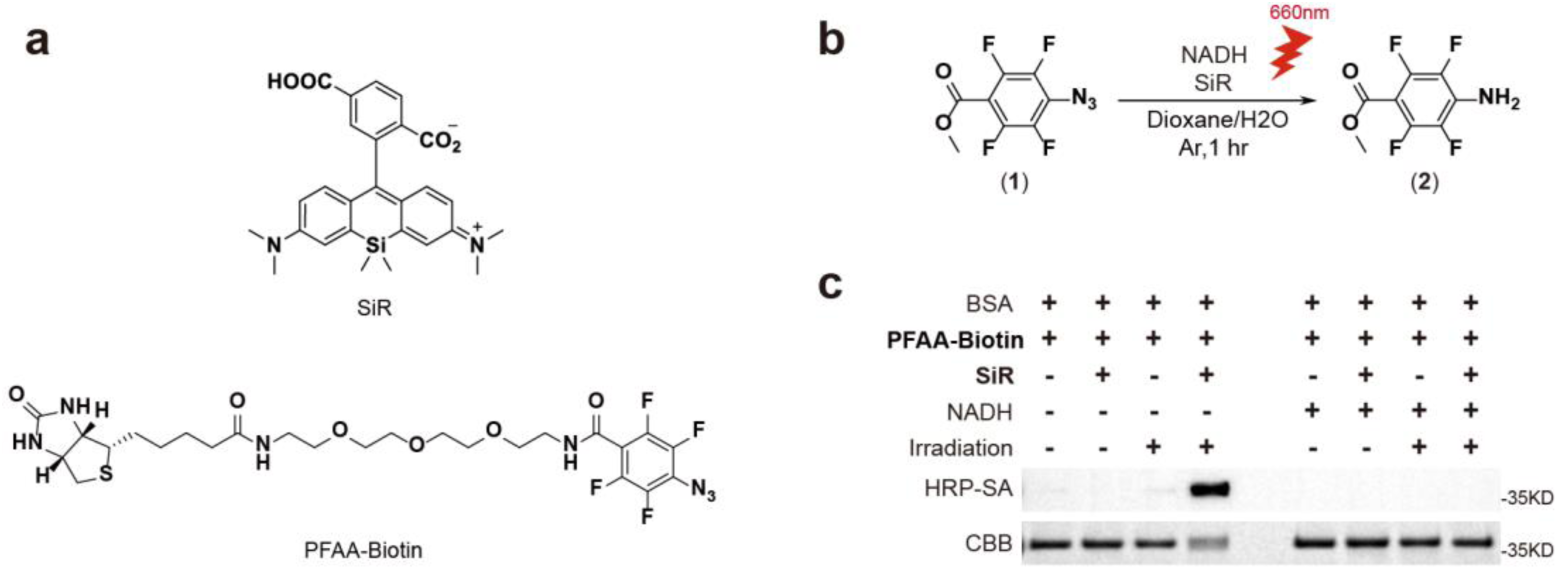
Test of electron transfer mediated proximity by SiR. **a,** Structures of photocatalyst and Biotin-conjugated probe. **b,** Reduction reaction of perfluorinated azide upon photoactivation. **c,** In vitro labeling of BSA. 0.2 μM BSA was mixed with 10 μM SiR, 100 μM PFAA-Biotin, and 2 mM NADH in PBS buffer. The mixtures were irradiated under 660 nm LED light of 200 mW/cm^2^ at 4 ℃ for 1 hour, followed by western blot to analyze the labeled result with HRP-SA.

**Extended Data Fig. 2.**
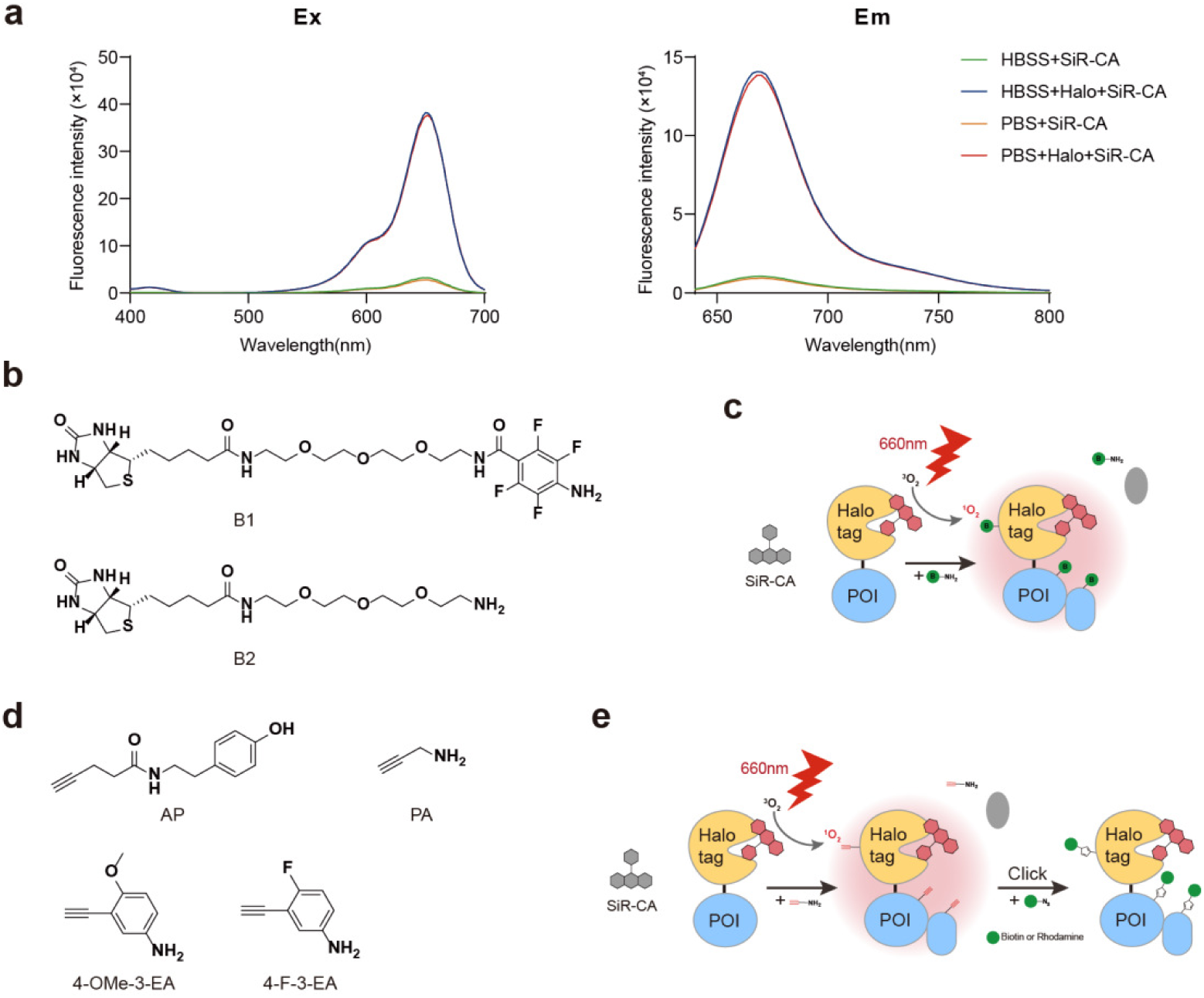
Photoactivated proximity labeling via the generation of singlet oxygen. **a,** Excitation and emission spectra of 10 μM SiR-CA in the presence or absence of 10 μM Halo protein in PBS or HBSS buffer. The excitation spectra were measured with emission at 750 nm and emission spectra were measured with excitation at 600 nm. **b,** The chemical structures of Biotin-conjugated probes. **c,** Schematic showing the Biotin-conjugated probe labeling. **d,** The chemical structures of alkynyl probes. **e,** Schematic showing the alkynyl probe labeling.

**Extended Data Fig. 3.**
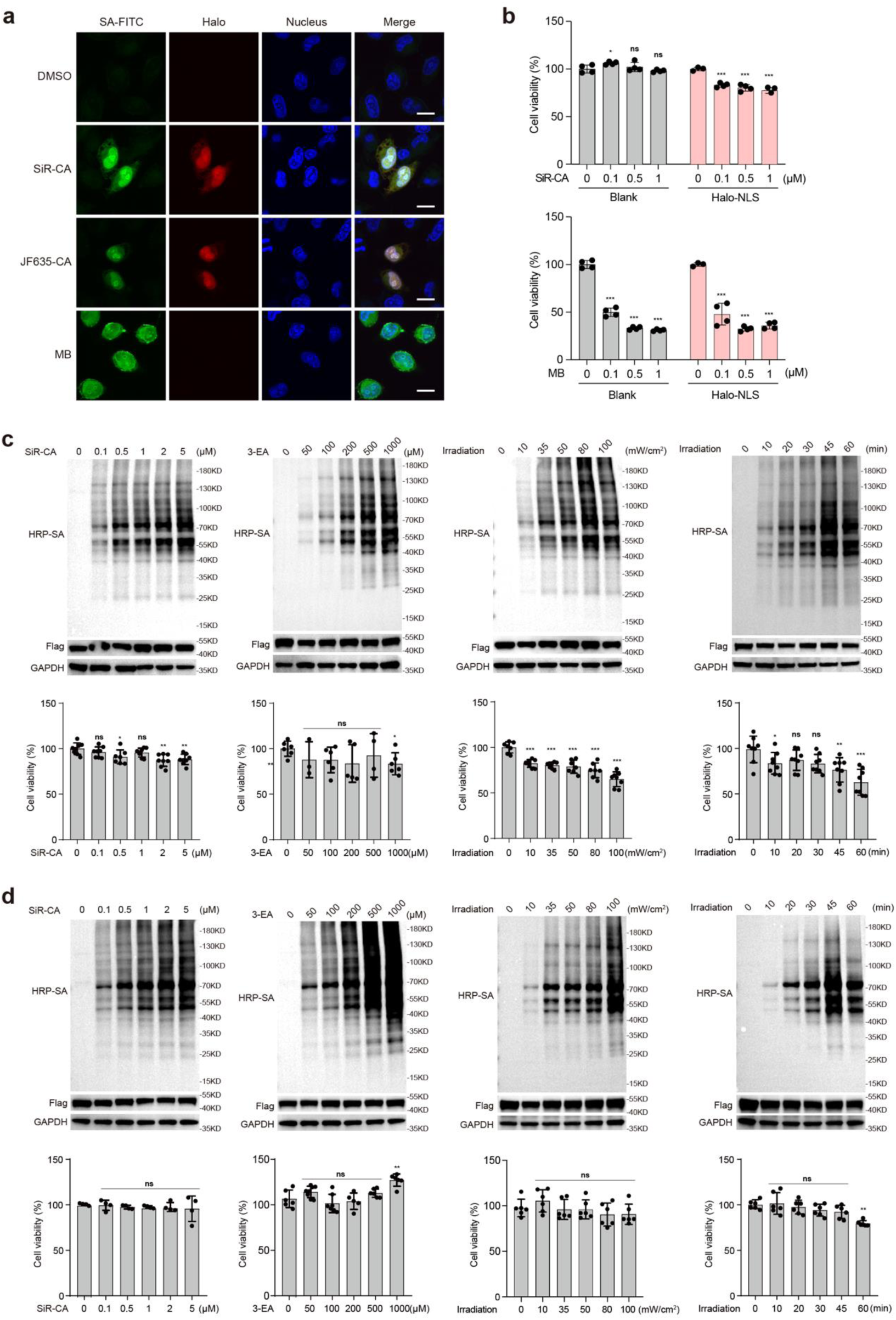
Validation and optimization of SeeID labeling system. **a**, Confocal images of HeLa expressing Halo-NLS labeled with different catalysts. Cells transiently expressing Halo-NLS were treated with 1 μM SiR-CA, JF635, or MB for 1 hour, and labeled by 3-EA under 660 nm light irradiation. Scale bars: 10 μm. **b**, Cytotoxicity of SiR-CA and MB. Blank and transiently expressing Halo-NLS HEK293T cells were treated with SiR-CA or MB for 1 hour, and irradiated under 660 nm LED light for 30 minutes. Phototoxicity was analyzed using Cell Counting-Lite 2.0. **c-d**, Optimization of SiR-CA concentration (with 500 μM 3-EA and irradiated at 50 mW/cm^2^ for 30 minutes), 3-EA concentration (with 1 μM SiR-CA and irradiated at 50 mW/cm^2^ for 30 minutes), irradiation intensity (with 1 μM SiR-CA, 500 μM 3-EA and irradiated for 30 minutes), and irradiation time (with 1 μM SiR-CA, 500 μM 3-EA and irradiated at 50 mW/cm^2^) in HeLa (**c**) and HEK293T (**d**). The cell viability was measured by Cell Counting-Lite 2.0.

**Extended Data Fig. 4.**
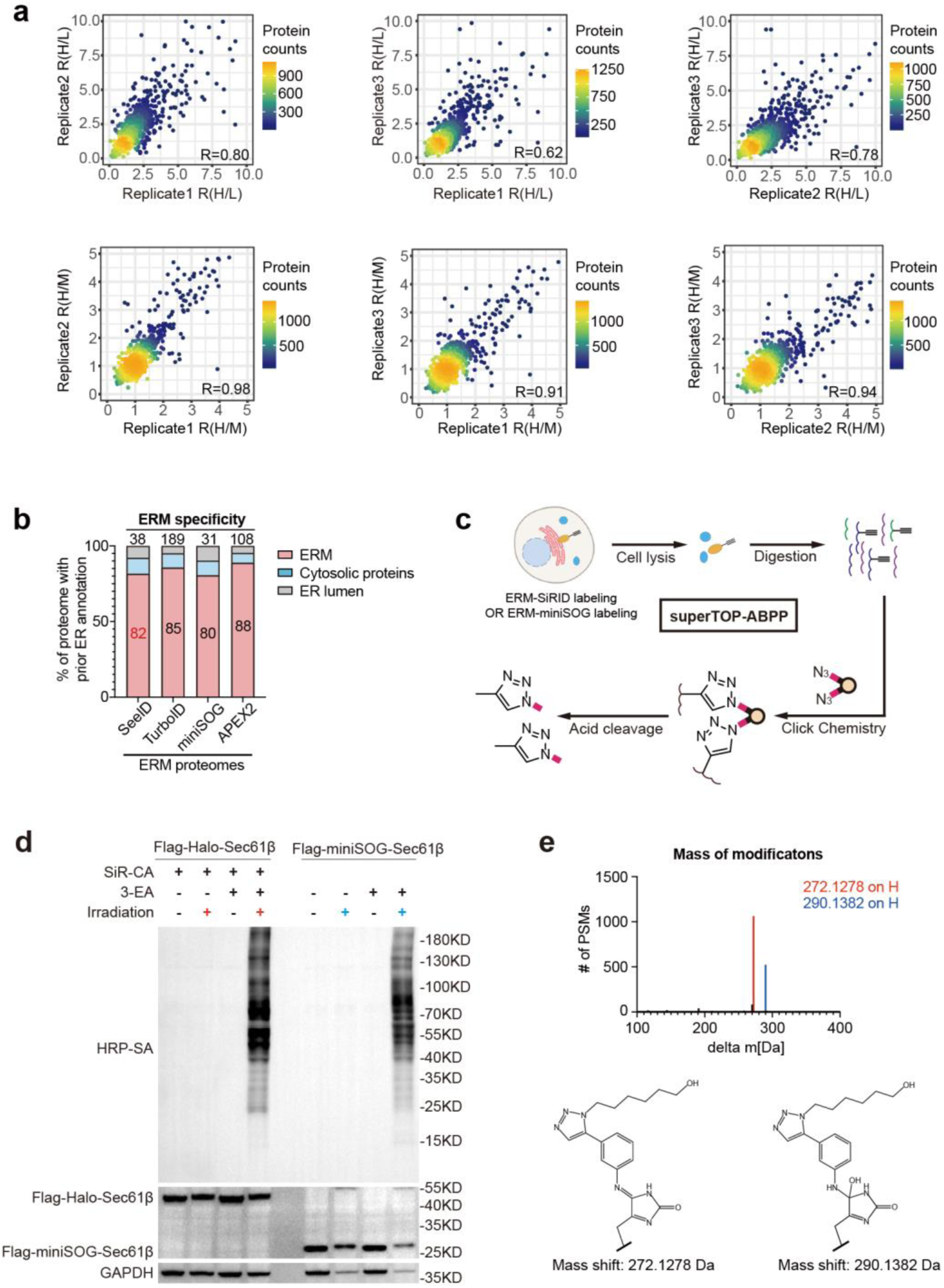
Profiling of endoplasmic reticulum proteome by SeeID. **a,** Volcano plot displaying the ERM data of **SeeID** labeling. **b,** Specificity comparison of **SeeID** with other proximity labeling methods at the ERM. **c,** Schematic of superTOP-ABPP to identify modification sites by **SeeID** and miniSOG labeling. **d, SeeID** demonstrated more efficient labeling than miniSOG. HEK293T cells transiently expressing Flag-Halo-Sec61β for 24 hours were treated with 1 μM SiR-CA and 500 μM 3-EA, and irradiated under 660 nm LED light for 30 minutes. Cells transiently expressing Flag-miniSOG-Sec61β for 24 hours were treated with 500 μM 3-EA, and irradiated under 450nm LED light for 20 minutes. **e,** SuperTOP-ABPP identified the number of modified histidine residues labeled by **SeeID** or miniSOG.

**Extended Data Fig. 5.**
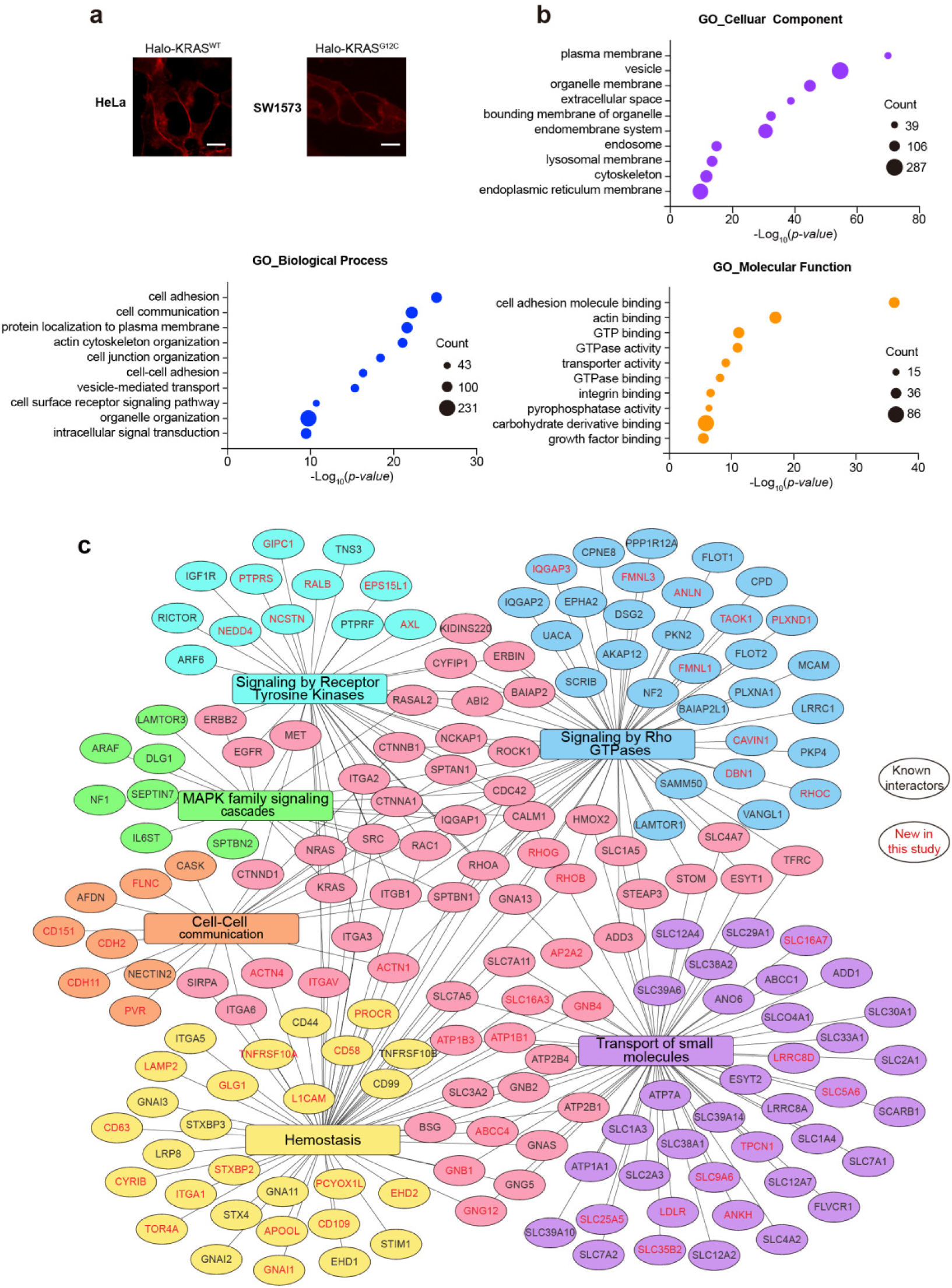
Profiling of KRAS proximity proteome by SeeID. a, Live cell imaging of Halo-KRAS^WT^ in HeLa and Halo-KRAS^G12C^ in SW1573. Scale bars: 10 μm. **b,** GO terms enriched for molecular functions, cellular components, and biological processes. **c,** Reactome pathway analysis of KRAS interactome identified by **SeeID**.

**Extended Data Fig. 6.**
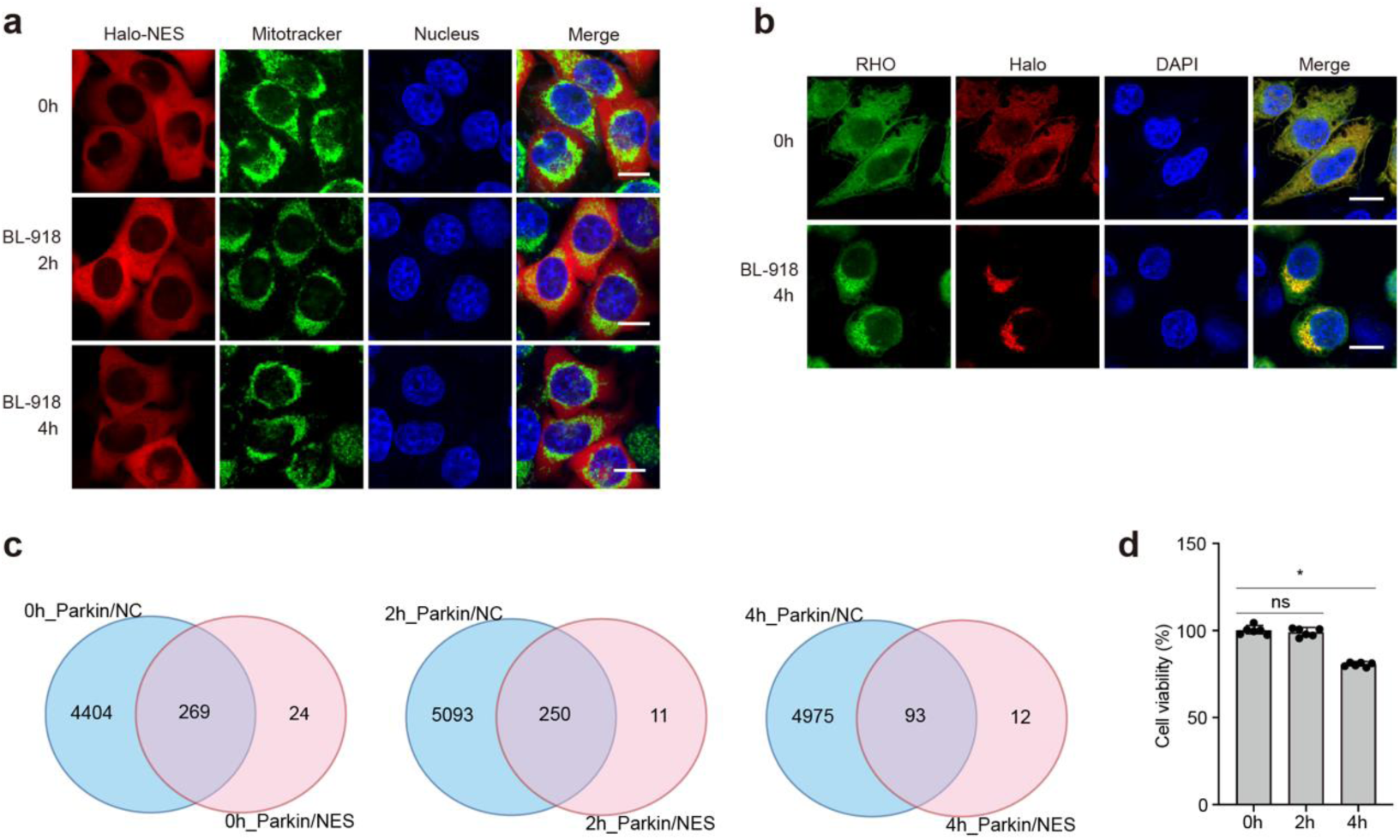
Imaging-guided profiling Parkin dynamic proteome. a, HeLa stably expressing Halo-NES were treated with 20 μM BL-918 for 0 hour, 2 hours and 4 hours. Mitotracker green were used as Mitochondria marker. Scale bars: 10 μm. **b,** Confocal immunofluorescence images of HeLa expressing of Halo-Parkin, cells were treated with BL-918 and subsequently labeled by **SeeID**. Scale bars: 10 μm. **c,** Overlapping of the identified proteins for indicated time. **d,** Cell viability after treatment of BL-918.

**Extended Data Fig. 7.**
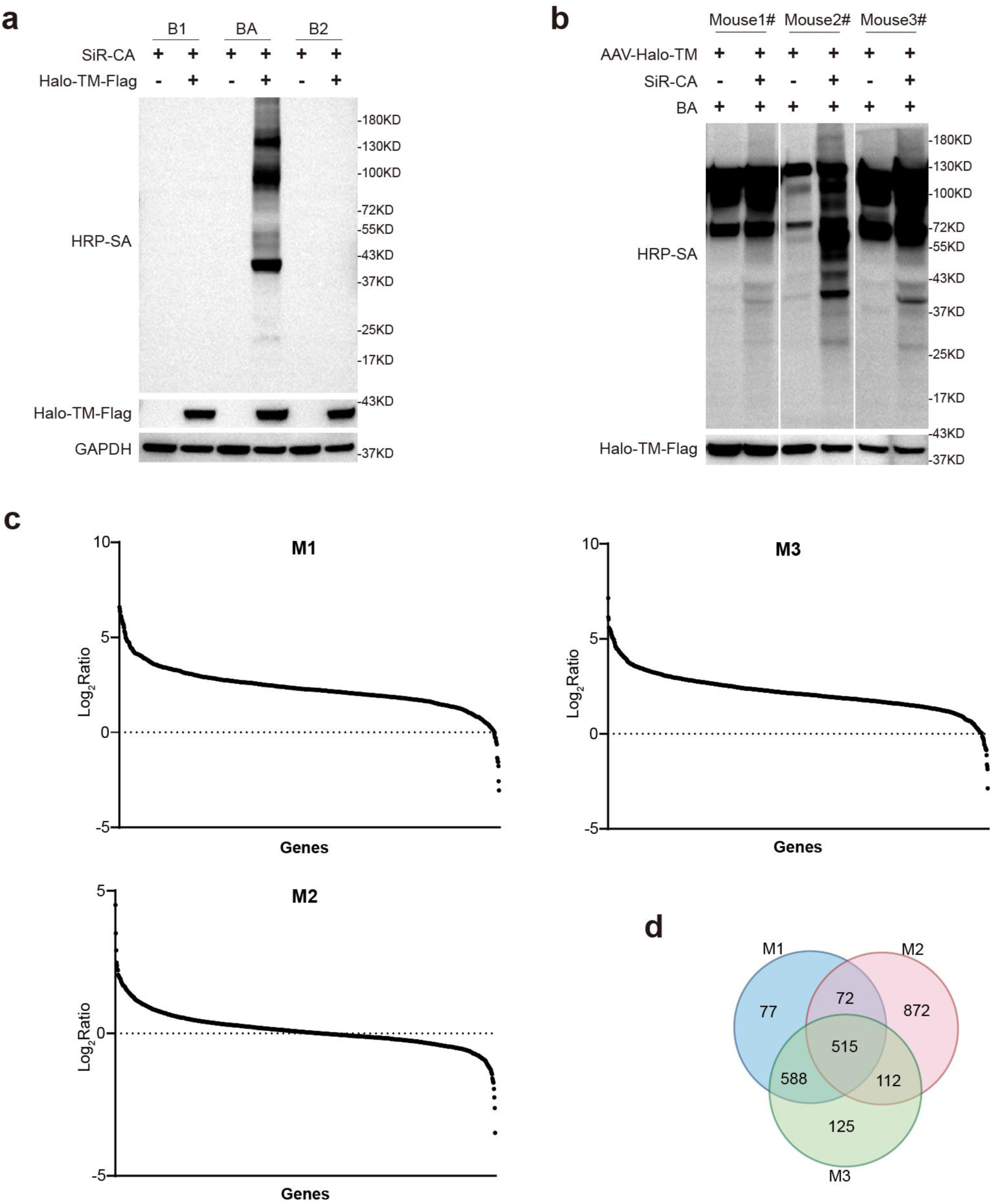
SeeID labeling in vivo. **a,** Investigation of the optimal Biotin-conjugated probe for plasma membrane protein labeling. HEK293T cells were transiently expressed Halo-TM-Flag for 24 hours, then treated with 1 μM SiR-CA for 1 hour and incubated with 500 μM probes for 10 minutes. Cells were irradiated under 660 nm LED for 30 minutes. **b,** Western blot analysis of cortex labeling. **c,** Log_2_Ratio(R/L) plot for the identified proteins in three biological replicates. **d,** Overlapping of enriched proteins in three mice.

